# A New Multiplex TaqMan qPCR for Precise Detection and Quantification of *Clavibacter michiganensis* in Seeds and Plant Tissue

**DOI:** 10.1101/2023.06.20.545733

**Authors:** Anne-Sophie Brochu, Tim J. Dumonceaux, Miryam Valenzuela, Richard Bélanger, Edel Pérez-López

## Abstract

Bacterial canker of tomato caused by *Clavibacter michiganensis* (*Cm*) is one of the most devastating bacterial diseases affecting the tomato industry worldwide. As the result of *Cm* colonization of the xylem, the susceptible host shows typical symptoms of wilt, marginal leaf necrosis, stem cankers, and ultimately plant death. However, is the ability of *Cm* to infect seeds and plants without causing symptoms what makes it an even more dangerous pathogen. Unfortunately, there are no resistant cultivars or effective chemical or biological control methods available to growers against *Cm*. Its control relies heavily on prevention. The implementation of a rapid and accurate detection tool is imperative to monitor the presence of *Cm* and prevent its spread. In this study, we developed a specific and sensitive multiplex TaqMan qPCR assay to detect *Cm* and distinguish it from related bacterial species that affect tomato plants. Two *Cm* chromosomal virulence-related genes, *rhu*M and *tom*A, were used as specific targets. The plant internal control *tubulin alpha-3* was included in each of the multiplexes to improve the reliability of the assay. Specificity was evaluated with 37 bacterial strains and more than 120 samples, including other *Clavibacter* spp. and related and unrelated bacterial pathogens from different geographic locations affecting a wide variety of hosts. Results showed that the assay was able to screen all *Cm* strains against other related bacteria. The assay was validated on tissue and seed samples following artificial infection and all tested samples accurately detected the presence of *Cm*. The tool described here is highly specific, sensitive, and reliable for the detection of *Cm* and allows the quantification of *Cm* in seeds, roots, stems, and leaves, finding a lower abundance of *Cm* in the roots compared to the other parts of the plant. The diagnostic assay can also be adapted for multiple purposes such as seed certification programs, surveillance, biosafety, the effectiveness of control methods, border protection, and epidemiological studies.

## INTRODUCTION

Bacterial canker of tomato is a systemic vascular disease caused by the gram-positive pathogen *Clavibacter michiganensis* (Smith, 1910) (*Cm*) (Davis et al. 1984; Li et al. 2018). It was first reported in the USA in 1909 (Smith 1910) and has been recognized as a devastating disease resulting in severe yield losses in tomato (*Solanum lycopersicum* L.) worldwide (Yang et Francis, 2005; Sen et al. 2015; Chalupowicz et al. 2017). The pathogen colonizes the xylem of the susceptible host causing wilt, marginal leaf necrosis, stem cankers, and ultimately plant death (Chalupowicz et al. 2012; Sen et al. 2015). The frequent occurrence of latent infections without visible symptoms (Chang et al. 1991; Gitaitis et al. 1991) and the ability of *Cm* to invade seeds (Franken et al. 1993; de León et al. 2006) make this pathogen particularly dangerous and easily dispersible. Currently, sanitation practices are the only available management strategy in the absence of resistant cultivars and effective chemical control measures (Sen et al. 2015; Nandi et al. 2018; Peritore-Galve et al. 2021). Pathogen-free seeds and transplants are essential for controlling the disease, while diagnostic methods are particularly needed to detect the presence of the bacterium and to manage the disease as fast as possible (Peritore-Galve et al. 2021). Several diagnostic methods have been developed for *Cm*, such as the use of semi-selective media followed by pathogenicity tests (EPPO, 2016; Ftayeh et al., 2011; Fatmi et al. 2017) and serological methods (Krämer et Griesbach, 1995; Kaneshiro et al., 2006). However, the methods developed so far are often unspecific, unreliable, and untimely (Dreier et al. 1997; Hadas et al. 2005; Kokošková et al. 2010; Sen et al. 2015).

In recent years, PCR-based molecular tools are the current choice for pathogen diagnosis due to their sensitivity, specificity, low cost, speed, and versatility (Mirmajlessi et al. 2015; Yang et al. 2021). Real-time PCR (qPCR) combines the advantages of PCR with quantitative measurement and a highly sensitive and specific tool for pathogen detection that can be used in routine diagnosis (Ouyang et al. 2013; Han et al. 2018; Ciampi-Guillardi et al. 2020; Ramachandran et al. 2021; Duong et al. 2022). It allows the detection of bacteria in an earlier stage of the infection, whether in a seed lot or asymptomatic plants. Multiplex qPCR can be used for increasing the specificity and the reliability of the detection by amplifying several targets of the pathogen and adding an internal control (Ramachandran et al. 2021; Ciampi-Guillardi et al. 2020).

Several genes have already been targeted to develop primer pairs specific to *Cm* (Dreier et al. 1997; Pastrik and Rainey 1999; Alvarez et al. 2005; Kaneshiro et al. 2006; Kleitman et al. 2008; Santos et al.1997; Ramachandran et al. 2021). However, new sequencing technologies have highlighted the genetic diversity present in the species (Thapa et al. 2017; Thapa et al. 2020; Ramachandran et al. 2021; Méndez et al. 2020). For example, the number of plasmids and their compositions can vary from one strain to another. Some strains have two plasmids (PCM1 and PCM2), while others have only one (Tancos et al. 2015; Croce et al. 2016; Thapa et al. 2017; Thapa et al. 2020; Méndez et al. 2020). In addition, there have been reported differences in the *chp*/*tomA* region among isolates from similar geographic regions, as seen in the case of Chilean isolates (Méndez et al. 2020). Furthermore, the recent identification of the presence of non-pathogenic *Clavibacter* in association with tomato seeds and plants has not been considered by current *Cm* detection tests, leaving room for unwanted false positive results (Jacques et al. 2012; Zaluga et al. 2013; Yasuhara-Bell and Alvarez 2015). Some primers previously developed target genes present in the plasmids or genes absent in some strains (Dreier et al. 1997; Kleiman et al. 2008), with several studies showing that most of the tests available can yield false negative and false positive results (Jacques et al. 2012; Thapa et al. 2020). Recently, a new multiplex PCR was developed by Thapa et al. (2020) using the virulence-related genes *rhu*M and *tom*A. This assay proved to be specific, sensitive, and reliable to detect pathogenic *Cm*, supporting their use in the development of a new quantitative diagnostic tool. In this study, we have designed and validated a new multiplex TaqMan qPCR assay for the detection of *Cm* in vegetative and seed samples that will contribute to the management of bacterial canker of tomato by detecting smaller amounts of bacteria faster and with higher specificity.

## METHODOLOGY

### Bacterial strains, growth conditions and DNA extraction

To validate our assay, 17 strains of *Cm* and 30 strains of genetically related bacteria species were used (Table 1). For several strains, we only had access to DNA, while others were generously donated by different collaborators (see acknowledgements), the provincial plant pathology diagnostic laboratory (*Laboratoire d’expertise et de diagnostic en phytoprotection*, MAPAQ) or isolated by us from infected tissues as previously described (Table 1; Ftayeh et al. 2011). Bacterial strains were grown in BD Bacto™ Tryptic Soy Broth (Soybean-Casein Digest Medium) at 28°C and 200 rpm overnight. All strains were stored at -80°C with a final concentration of 20% glycerol, while DNA was conserved at -20°C until analysis. Genomic DNA was extracted from bacterial cultures using the cetyltrimethylammonium bromide (CTAB) method as previously described (Murray and Thompson 1980).

**Table 1.**
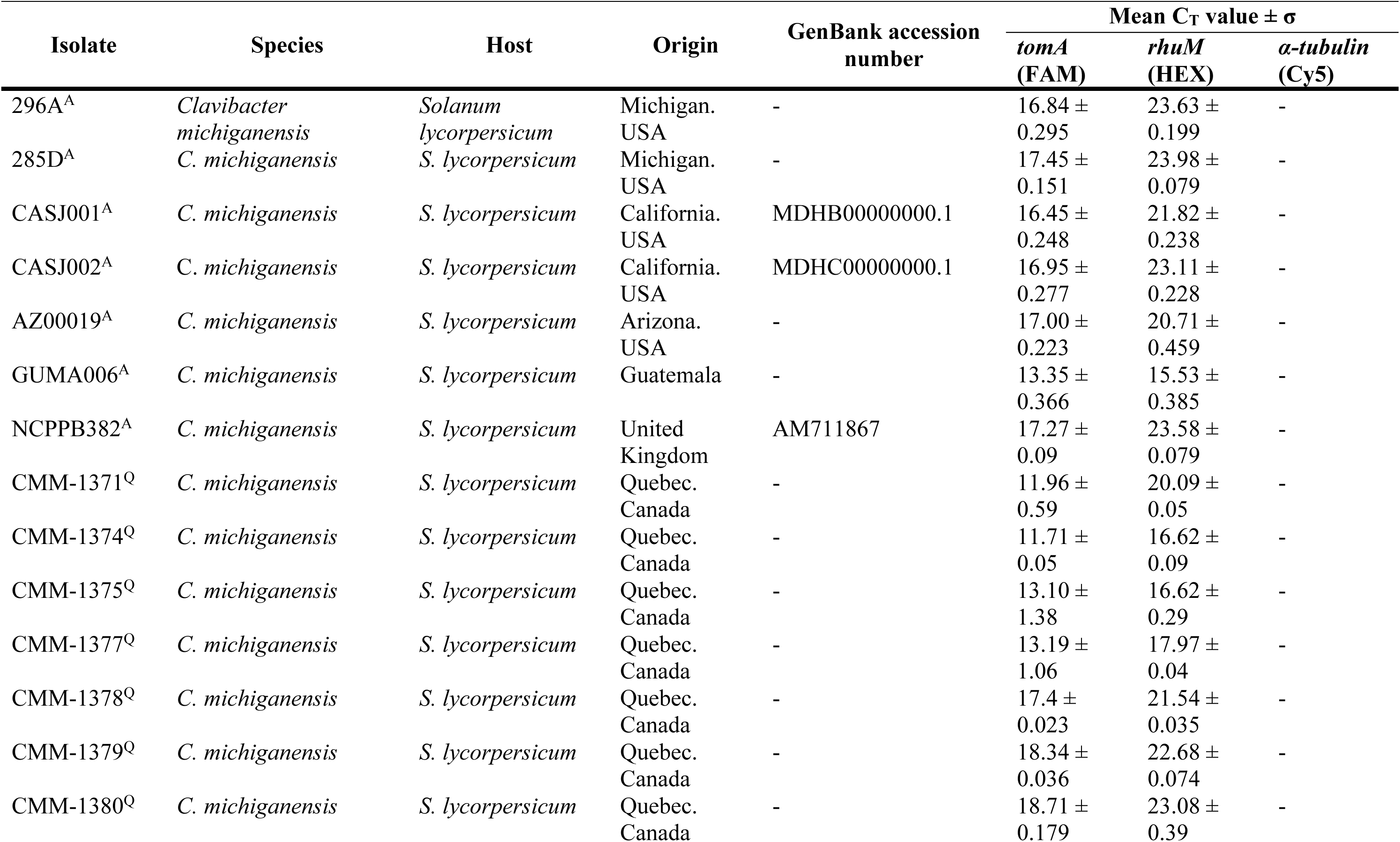
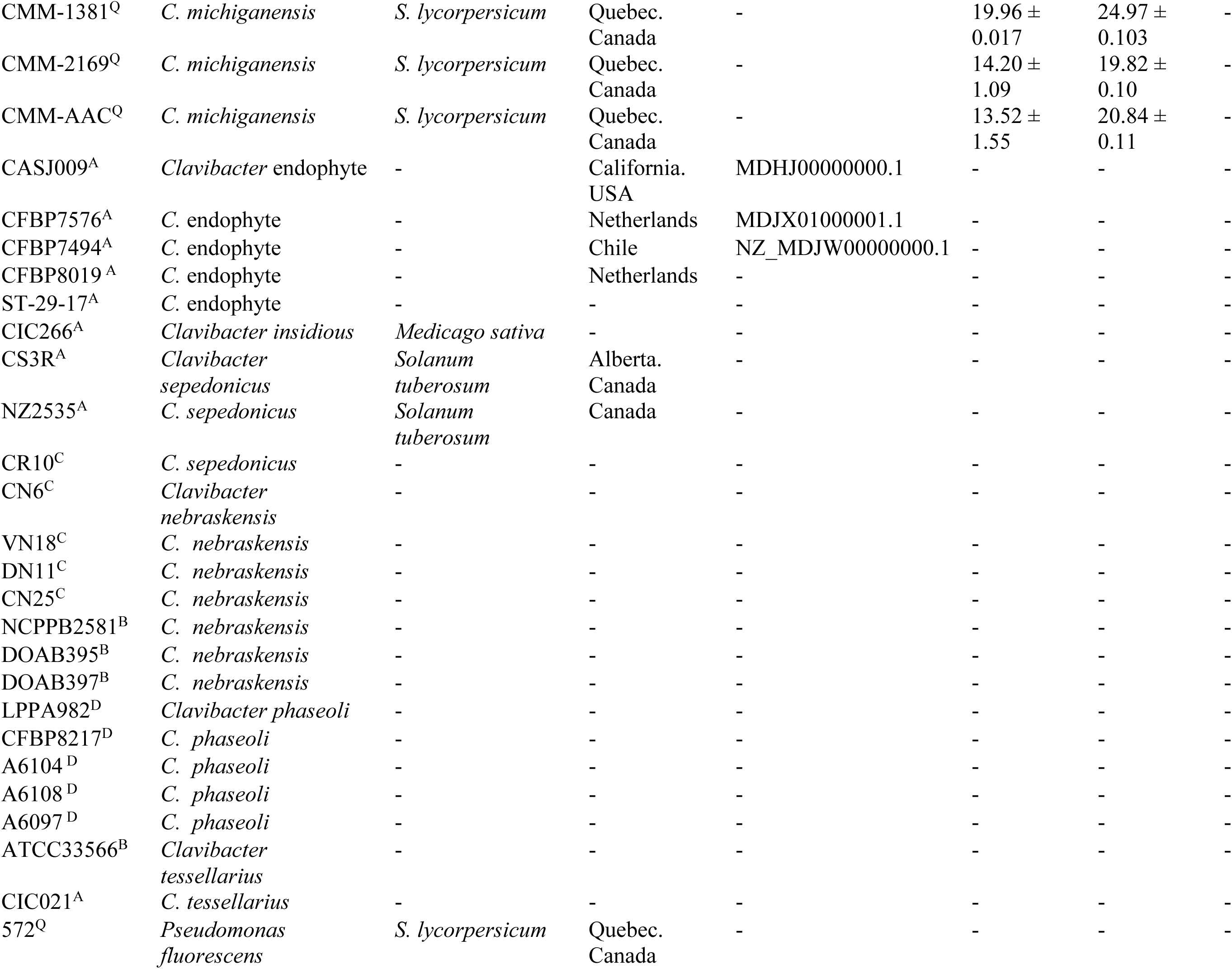
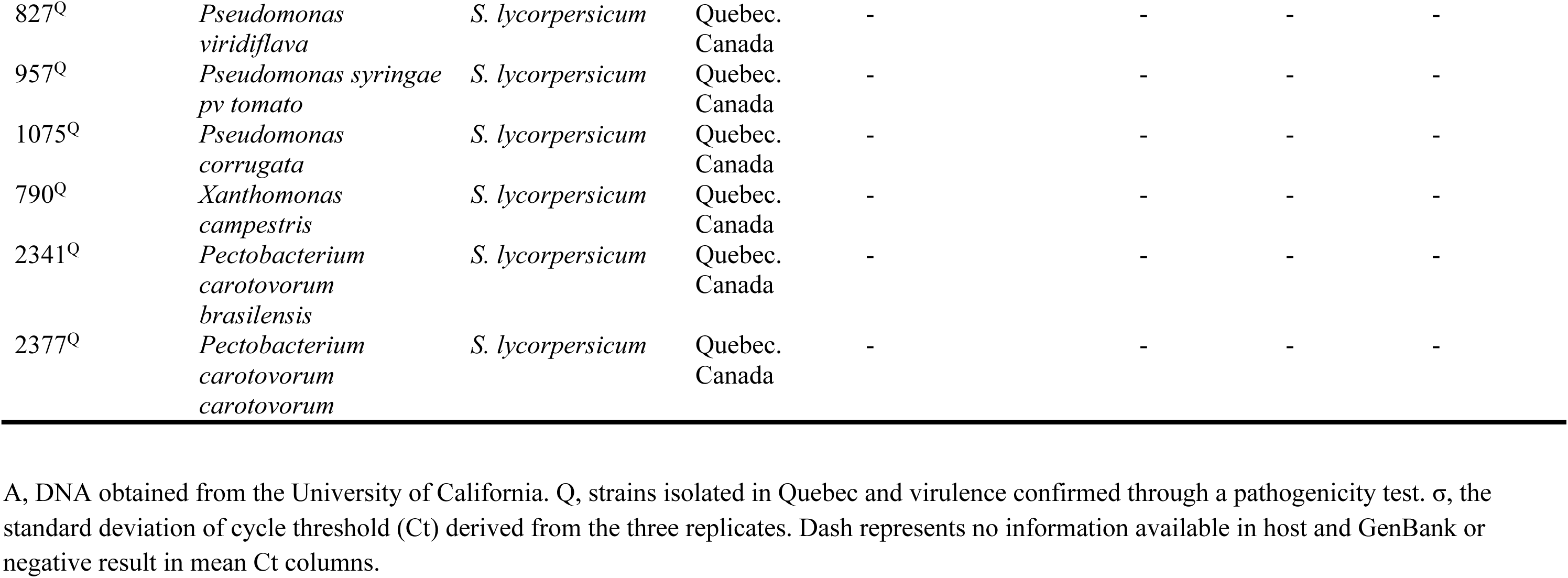
Isolates used in this study to validate the TaqMan multiplex qPCR.

### Bacterial strains validation

Bacterial strains isolated from infected tissues were validated using a pathogenicity test on the tomato cultivar M82. Plants were grown under greenhouse conditions at 26°C during the day and 18°C at night with 60% relative humidity and a 16-hour photoperiod. Tomato plants at the 2- to 3-leaf stages were inoculated with different isolates. Plants were wounded at the cotyledons and inoculated, using a syringe, with 10 µl of bacterial solution. The bacterial solution was diluted with 10 mM MgCl_2_ to an OD_600_ of 0.1, approximately 10^8^ CFU/ml (Epoch 2 Microplate Spectrophotometer). Tomato plants used as control were inoculated with 10 mM MgCl_2_. Symptoms were recorded after 7–21 days. For each strain, five plants were inoculated. To confirm the presence of pathogenic *Cm* in the plant, we used the multiplex PCR assay as previously described (Thapa et al. 2020)

### Primer and probe design

Specific primers and hydrolysis probes were designed for the detection of *Cm* using as templates the sequence of the tomatinase (*tomA*) gene, the *chp/tomA* genomic island gene, and the hypothetical protein *rhuM,* previously amplified, sequenced and deposited into GenBank (NC_009480.1) (Thapa et al. 2020). A third set of primers and probe was designed to amplify the tomato *tubulin alpha-3* chain (GenBank accession no. LOC101254013) as an internal control for the multiplex. All primers and probes were designed using the Beacon Designer software v8.0 and synthesized by Integrated DNA Technologies Inc. Each primer set was designed to have similar melting temperatures with 8 to 10 °C lower than the probes and to amplify products of less than 200 bp (Table 2). Thermodynamic features, self-dimer occurrence and the presence of secondary structures were analyzed *in silico* using the web server OligoAnalyzer (https://www.idtdna.com/pages/tools/oligoanalyzer). Primer and probe specificity was analyzed using the primer designing online tool Primer-Blast (https://www.ncbi.nlm.nih.gov/tools/primer-blast/).

**Table 2:**
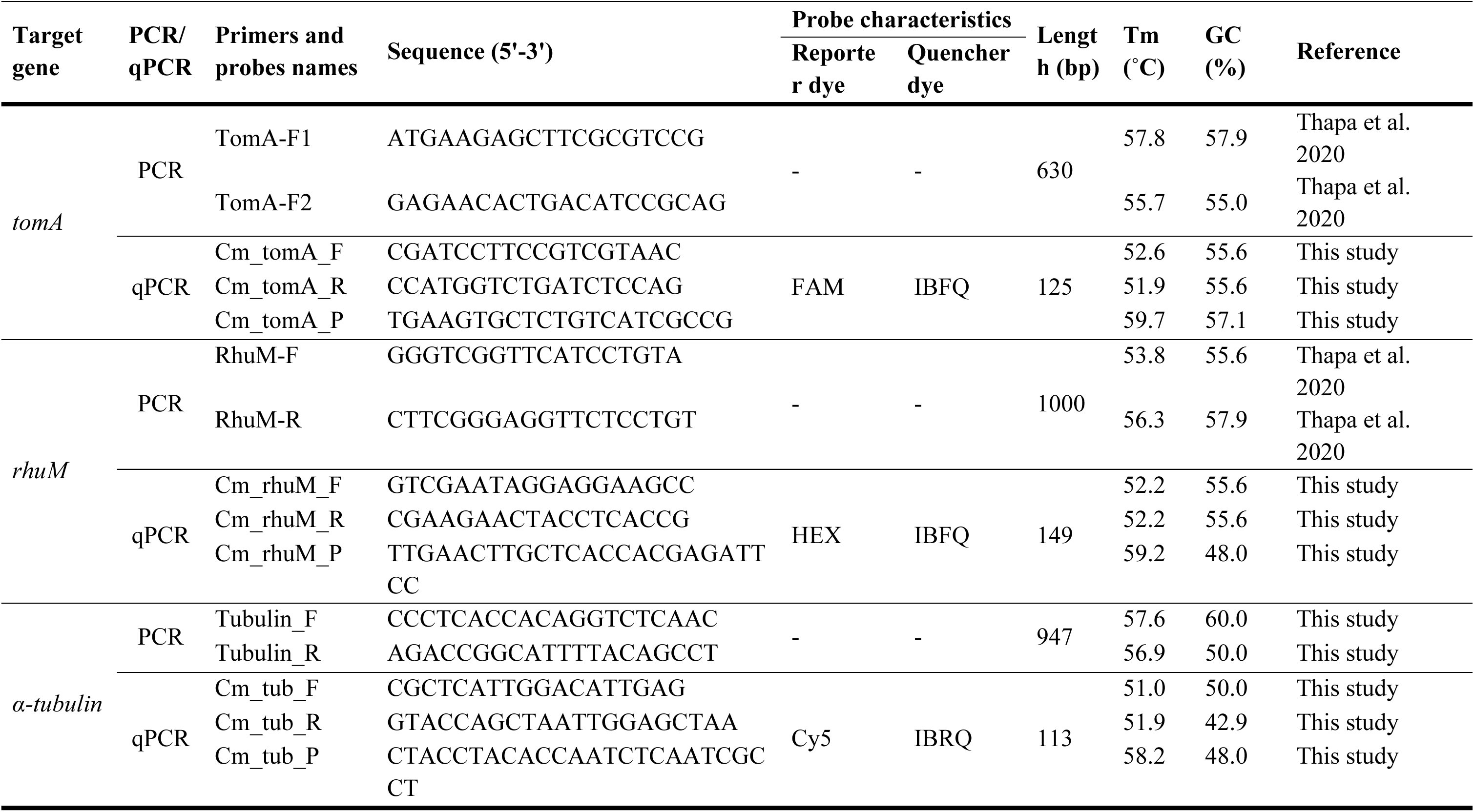
Information and characteristics of primers and probes designed or used in this study for the specific detection of *Clavibacter michiganensis*.

### TaqMan multiplex and single qPCR conditions

Detection of bacterial DNA in the different samples was performed with TaqMan technology using the CFX Opus Real-Time PCR Systems (Bio-Rad). Multiplex TaqMan qPCR assays were prepared in 20 µl of reaction mixtures containing 1X iQ^TM^ Multiplex Powermix (Bio-Rad, CAD), 300 nM of the sense and antisense primers for each target, 200 nM of each probe and 2 µl of DNA template. The volume was adjusted with DNA- and RNA-free water (Bio-Rad, CAD). Thermal cycling conditions were 95 °C, 3 min (1×); 95 °C, 10 sec/60 °C, 30 sec (40×); and data collection. Emitted fluorescence was calculated after each extension for all channels. Each sample was analyzed in triplicate. In all analyses, DNA- and RNA-free water was included as non-template control (NTC). TaqMan qPCR assays for single markers were performed similarly but for each set of primers/probes independently using 1X SsoAdvanced™ Universal Probes Supermix (Bio-Rad, CAD). Thermal cycling conditions were 95 °C, 3 min (1×); 98°C, 15 sec/60°C, 45 sec (40×), and data collection. Emitted fluorescence was calculated after each extension for all channels. Each sample was analyzed in triplicate. The experimental data obtained were analyzed with CFX Maestro Software 2.3 (Bio-Rad, CAD).

### Specificity of the primers

The specificity of the primers was first evaluated using SYBR green qPCR in 20 µl reactions containing 1X iQ™ SYBR® Green Supermix (Bio-Rad), 300 nM of each primer, 2 µl of DNA template and DNA- and RNA-free water up to the final volume. Thermal cycling conditions were 95°C, 3 min (1×); 95°C, 10 sec/60°C, 45 sec (40×), and data collection. Furthermore, the melt curve and peak analyses were carried out immediately at a melting rate value of 0.5 °C/5 s, from 65 to 95 °C. Each sample was analyzed in triplicate. The experimental data were analyzed using CFX Maestro Software 2.3 (Bio-Rad, CAD). Genomic DNA from healthy plants, *Cm* infected plants, *Cm tom*A*, Cm rhu*M and *tomato tubulin alpha 3* plasmids were generated as described below, and no template control was used to analyze each primer pair. The amplification product was analyzed through electrophoresis using 1.5% agarose gel stained with SYBR Safe (ThermoFisher, CAD) and visualized under UV light.

### Multiplex TaqMan qPCR assay specificity

The specificity of the multiplex qPCR assay was evaluated using genomic DNA isolated from 17 *Cm* strains and 30 strains of genetically related bacteria species (Table 1). Genomic DNA extracted from the stem tissue of healthy tomato plants and healthy tomato seeds was also included to validate cross-reactions of the designed primers (Table 2). Genomic DNA from *Cm* isolate Cmm-1375 (Table 1) was used as a positive control in each assay, along with no template control. Conditions and data analysis were performed as described above. Each sample was analyzed in triplicate. The same samples were analyzed using the multiplex PCR assay previously described by Thapa et al. (2020) to detect pathogenic *Cm* used here as the ‘gold standard’.

### Multiplex TaqMan qPCR assay sensitivity

The sensitivity of the multiplex qPCR assay was evaluated using a serial dilution of plasmids containing the genes of interest. The genes were partially amplified through PCR with the primers presented in Table 2. PCR was performed in 50 μl of final volume with 2 μl of DNA; 0.5 μM forward and reverse primers, 1 X Phusion HF buffer, and Phusion DNA Polymerase (New England Biolabs, CAD) at a final concentration of 0.02 U/μl. All PCR amplifications were carried out in a T100 Thermal Cycler (Bio-Rad). Thermal cycling conditions for *tom*A*, rhu*M and tomato *tubulin alpha-3* were respectively 98 °C, 30 sec (1×); 98 °C, 10 sec/63.6°C (*tom*A), 61.3 °C (*rhu*M), 65 °C (*tomato tubulin alpha-3)*, 30 sec/72 °C, 30 sec (35×); 72 °C, 10 min. PCR products were observed through electrophoresis using 1% agarose gels. Amplicons of each gene were cloned using the pGEM®-T Vector System (Promega, CAD) into competent *E. coli* Top10 cells according to the manufacturer’s instructions. Clones carrying specific plasmids were selected using LB Broth, Miller (Luria-Bertani) containing 100 µg/ml of ampicillin (Sigma Aldrich, CAD) at 37°C. Plasmids were purified using GeneJET Plasmid Miniprep Kit (Thermo Scientific, CAD) according to the manufacturer’s instructions and the presence of the expected sequence was confirmed through restriction analysis and sequencing using plasmid-targeted M13 sequencing primers forward and reverse (Centre Hospitalier de l’Université Laval de Quebec, Canada).

To prepare standard curves, plasmids were linearized using SpeI restriction endonuclease following the manufacturer’s recommendation (New England Biolabs, CAD). The concentration of the linearized plasmid DNA was quantified using the Qubit 4 fluorometer (Thermo Fisher, CAD), and *Cm tomA, Cm rhu*M and *tomato tubulin alpha-3* linearized plasmids were serially diluted with DNA- and RNA-free water (Bio-Rad, CAD) and stored at −20°C. The number of copies/μl was calculated as (NA × C)/MW, where NA is the Avogadro constant expressed in mol^−1^, C is the concentration expressed in g/μl, and MW is the molecular weight expressed in g/mol. Efficiency, slope and linear correlation of primer and probe pairs were determined using CFX Maestro Software 2.3 (Bio-Rad, CAD). Limits of quantification (LOQ) and detection (LOD) were calculated for the qPCR assay using the LoD-calculator R script by Klymus et al. (2019). This script uses a curve-fitting approach based on the R drc package, running all available logarithmic functions, and selecting the best-fitting model using the mselect function in the drc package. Best-fitting models from the LoD-calculator script were used to report modelled LOQ and LOD, with a 95% LOD probability of detection and a 35% CV for LOQ precision.

The sensitivity of the multiplex TaqMan qPCR was also evaluated using Cmm-1375 genomic DNA. Genomic DNA was extracted from an overnight bacterial culture at 4 × 10^9^ CFU ml^−1^ using a CTAB method. Bacterial concentration was estimated by plating serial dilutions on TSA medium and incubating the plates for 24 h at 28°C and counting the colonies at each dilution. Then, the genomic DNA was used to prepare a standard curve by serially diluting 10^8^ copies of genomic DNA 10-fold from 10^7^ to 10 copies.

### Validation of the assay

#### Plant samples

Twelve tomato plants were inoculated as described above with the *Cm* isolate Cmm-1375 and three others were inoculated with 10 mM MgCl_2_. Samples were harvested 21 days post-inoculation (dpi). To obtain DNA from inoculated plants, stem fragments (1 cm each) were collected at the inoculation site, 3 and 6 cm above the inoculation site, from the rachis of the third leaf and the roots. The samples were dried using a freeze dryer and macerated in a 2-ml tube. Then, grounded tissue was used for DNA extraction using a CTAB method. Conditions and data analysis were performed as mentioned above.

Stem fragments (1 cm each) were collected at the inoculation site, 3 and 6 cm above the inoculation site, from the rachis of the third leaf and the roots. Two 2.8 mm ceramic bulk beads (Omni International, CAD) and 500 µl of sterile ddH_2_0 were added to the samples in 2 ml tubes. The tissue was ground using Bead ruptor elite (Omni International, CAD) at two cycles 40 seconds at 5 m/s with a pause of 40 seconds in between the two cycles and 100 µl of the macerate was added and spread onto TSA solid media. After 24 h at 28 °C, several colonies were selected to do colony PCR using the multiplex test as described above.

#### Statistical analysis

The effects of standard curves, genes, and tissues on the number of *Cm* copies g^-1^ were studied using a linear mixed model on the data since the response variable is continuous. In this model, standard curves (2 levels), genes (2 levels), and tissues (5 levels), as well as all two- and three-way interactions were considered as fixed factors, whereas plants were used as random factors. The analysis was performed by transforming the number of *Cm* copies g^-1^ measures by the logarithm, as suggested by the Box-Cox technique, to satisfy the normality assumption. Following a significant effect of one source of variation, we used Tukey’s multiple comparisons method to identify the levels of that source that differed from the others, while controlling for the type I error rate. All analyses were performed using R software version 4.2.3 (R Core Team, 2023) with the nlme package lme function for the linear mixed model, at the 0.05 significance level. The code and raw data used in this study can be found using this link https://github.com/Edelab/Development-of-a-multiplex-TaqMan-qPCR-targeting-Clavibacter-michiganensis-virulence-related-genes.

#### Seed samples

For seed contamination, the method developed by Thapa et al. (2020) was modified to suit our laboratory accommodations. The inoculum was prepared by diluting *Cm* isolate Cmm-1375 with 10 mM MgCl_2_ to an OD_600_ of 0.1 (Epoch 2 Microplate Spectrophotometer). Lots of 1 g of tomato seeds were incubated with 4 mL of Cmm-1375 bacterial solution for 24 h at 37°C. After 24 h, the liquid was removed, and the seeds were dried at 24°C on filter paper for 2 days and stored for 4 weeks at room temperature. To evaluate the detection limit, we generated and tested three different infested: healthy seed mixtures in the following ratios, (*i*) 1:399, (*ii*) 1:799, and (*iii*) 1:1599, which corresponds to infestation levels of 0.25, 0.125, and 0.0625%, respectively. Seed mixtures were resuspended in seed extraction buffer pH 7.4 (7.75 g of Na_2_HPO_4_, 1.65 g of KH_2_PO_4_ and 0.5 mL of Tween 20 for a litre). Samples were incubated at 4 °C for 14 h and later transferred to a heated bath and incubated at 65 °C for 30 min, vortexing each 5 min. The liquid was pipetted out of the mix transferred to a 2-ml tube for DNA extraction tube and centrifuged for 3 min at 14 000 g. The supernatant was discarded, and DNA was extracted from the pellet using a CTAB method. Conditions and data analysis were performed as mentioned above. The results were compared with the multiplex PCR assay to detect *Cm* in seed as described previously (Thapa et al. 2020). The detailed protocol followed was added as supplementary.

## RESULTS

### Primer and probe specificity

*In silico* analysis of the sets of primers and probes targeting *tom*A and *rhu*M using BLASTn with the NCBI GenBank database, revealed 100% nucleotide identity over 100% query cover with *Clavibacter michiganensis*. No nucleotide similarity was recorded for tomato, other *Clavibacter* species or other species. These results indicate that *tom*A and *rhu*M are highly conserved among *Cm* strains and absent or highly diverse in other bacteria. Primers designed to amplify tomato *tubulin alpha-3* used here as control did not hybridize with *Cm* nucleotide sequences available in the NCBI GenBank databases.

The specificity of each primer pair was evaluated through SYBR green qPCR with genomic DNA from healthy plants, *Cm* infected plants, *Cm tom*A*, Cm rhu*M and tomato *tubulin alpha-3* plasmids, and no template control (Fig. 1). For *tom*A and *rhu*M genes, we verified the presence of only one PCR product, for the specific plasmids and infected plants corroborated by a unique melting peak at Tm = 86.75°C and Tm = 89°C, respectively (Fig. 1A-B). Agarose gel electrophoresis verified the expected product size of 125 bp for *tom*A and 149 bp for *rhu*M with no amplification in the negative control and the healthy plant (Fig. 1D). In the case of the SYBR Green qPCR for tomato *tubulin alpha-3* gene amplification, we observed for the specific plasmids, infected plant samples and healthy plants, a very clear peak at Tm = 79°C (Fig. 1C), confirmed by the presence of only one amplification product of the expected 113 bp in an agarose gel (Fig. 1D).

**Figure 1.**
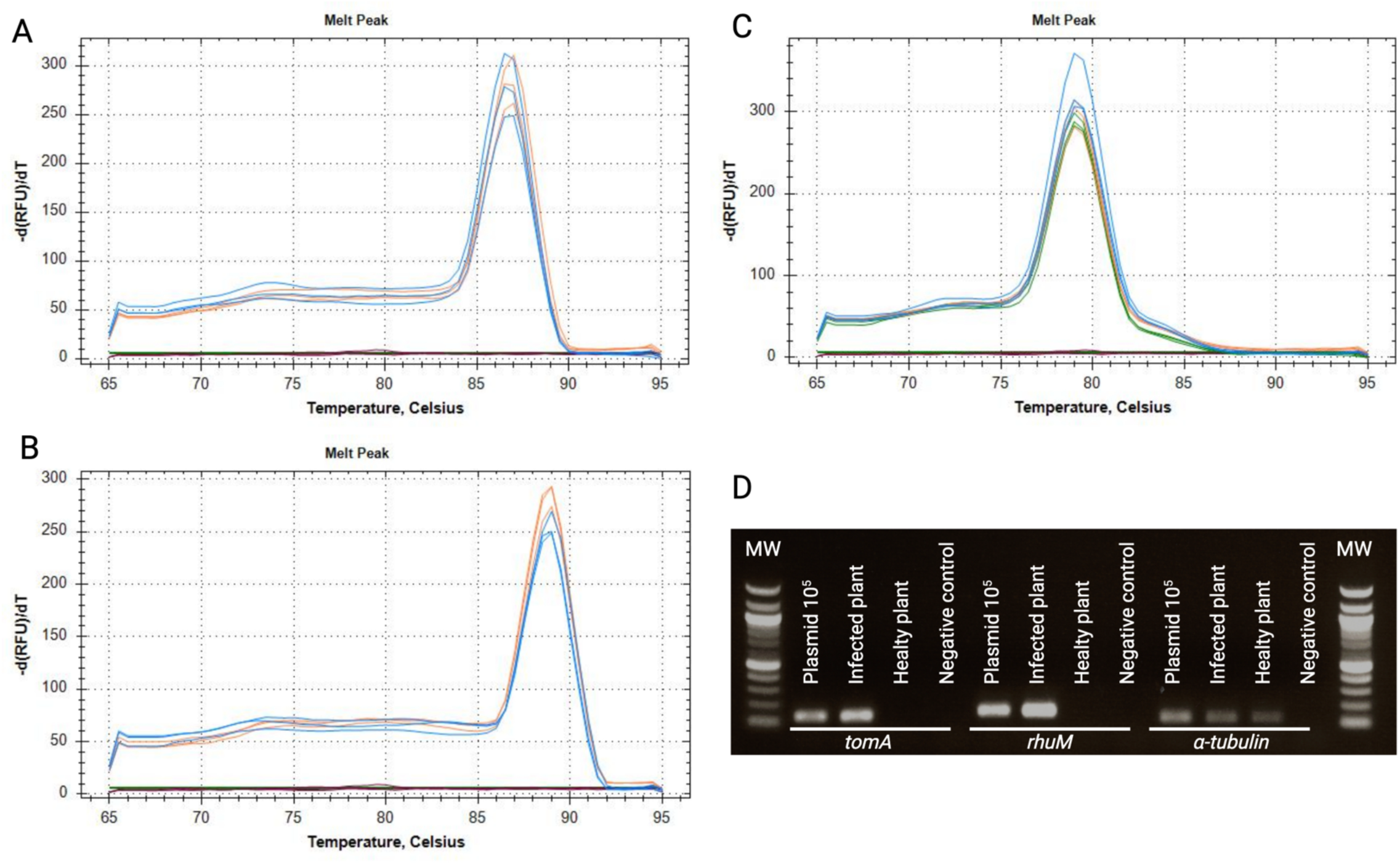
Specificity of primers and probes using SYBR Green qPCR. **A,** *tom*A melting peaks. **B**, *rhu*M melting peaks. **C**, *α-tubulin* melting peaks. In blue, plasmids containing each gene; orange, genomic DNA from *Clavibacter michiganensis*-infected tomato plants; green, genomic DNA from healthy tomato plants; red, no template control. **D**, Agarose gel electrophoresis of SYBR Green qPCR products. MW, molecular weight marker 100 bp DNA ladder.

### Specificity of multiplex TaqMan qPCR assay

All *Cm* strains tested here were detected with high accuracy and specificity using primers *Cm_ tom*A *_*F/*Cm_ tom*A *_*R and probe *Cm_ tom*A *_*P, primers *Cm_rhuM_*F*/Cm_rhuM_*R and probe *Cm_rhuM_*P (Table 1). Primers *Cm_tub_*F*/Cm_tub_*R and probe *Cm_Tub_*P did not amplify as expected when using bacterial genomic DNA (Table 1). In all qPCR assays performed, strain Cmm-1375, used as positive control, amplified for both primer/probe sets except for the tomato *tubulin alpha-3* gene and no cross-reactivity with any other target was detected, while no amplification was observed in the NTC. The three fragments, *rhu*M (1 000 bp), *tom*A (630 bp), and 16S rRNA (415 bp) were amplified from *Cm* strains, while only the fragment corresponding to 16S rRNA was amplified from other bacteria (Fig. 2).

**Figure 2.**
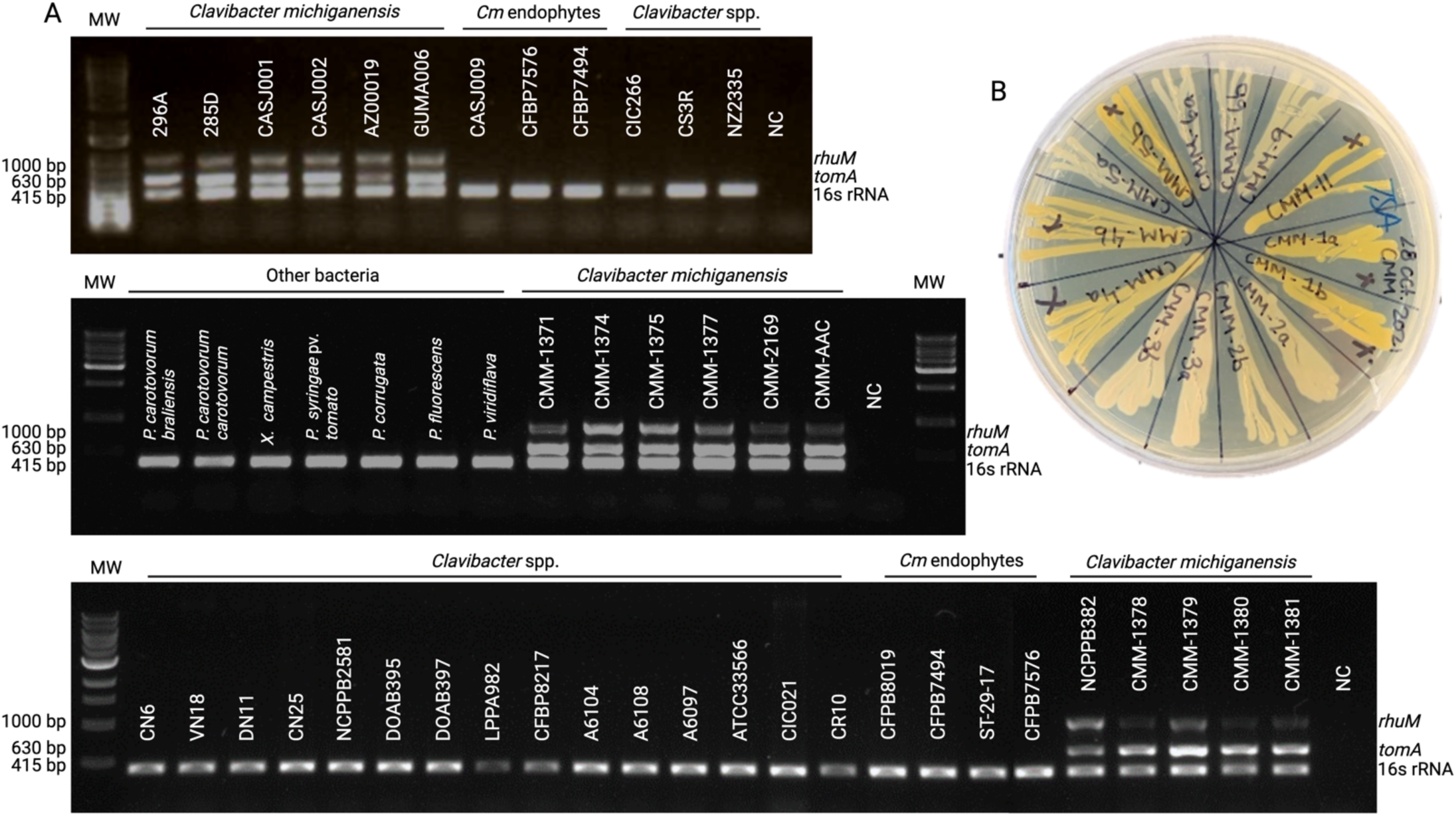
Characterization of *Clavibacter michiganensis* and bacterial strains used in this study with the multiplex PCR developed by Thapa et al (2020). Detailed information of the strains can be found in Table 1. A, Agarose gel electrophoresis, MW, molecular weight marker 100 bp DNA ladder. B, representation of the *Cm* strains used in this study.

### Sensitivity and efficiency of multiplex TaqMan qPCR assay

The limit of detection (LOD) of the TaqMan qPCR multiplex was determined by performing a serial dilution of each plasmid containing the genes of interest. The assay showed a LOD of 10 copies per reaction when using purified plasmids for all targets (Fig. 3), and 100 copies when using pure *Cm* bacterial culture (Fig. 3). In addition, results were obtained for the plasmid DNA assay using an optimized curve-fitting model, 95% LOD of 50 copies per reaction for the *tom*A and *rhu*M genes whereas 1000 copies per reaction for tomato *tubulin alpha-3* (Supplementary Table S1).

**Figure 3.**
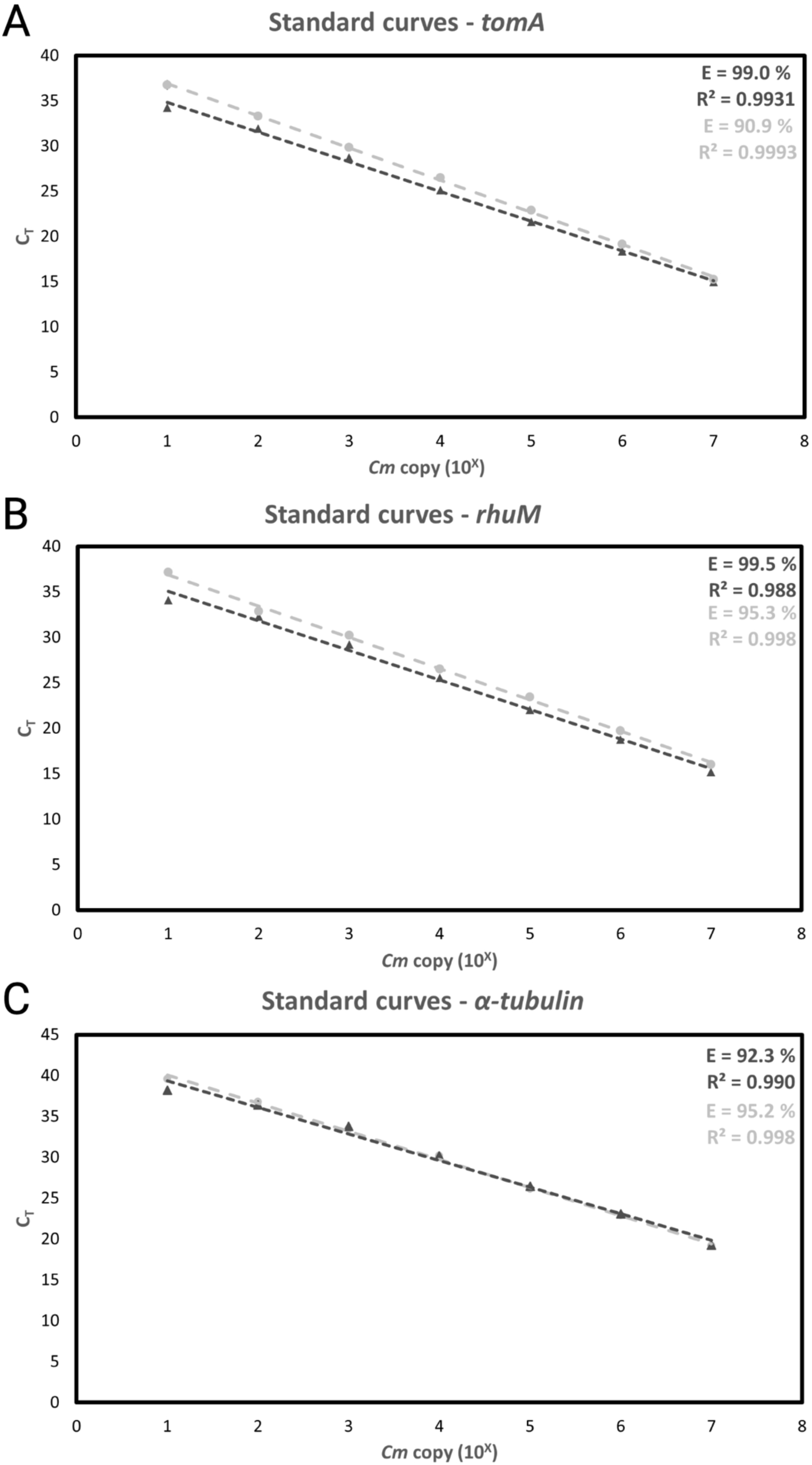
Standard curves for the TaqMan qPCR designed in this study using seven-fold serial dilutions of plasmids containing three targets: *tom*A (A), *rhu*M (B), and *α-tubulin* (C). The number of copies ranged from 10^7^ to 10^1^ copies. Dot/light gray represents single-plex TaqMan qPCR assays and triangle/dark gray represents the results in multiplex TaqMan qPCR. The values presented in the graph represent the mean Ct of three technical replicates.

For the purified plasmid DNA and pure *Cm* bacterial culture sensitivity assays, statistical parameters such as reaction efficiency, slope and linear correlation for primer sets and probes targeting *Cm* and tomatoes were in an optimal range, with efficiency between 90% and 110% and R^2^ between 0.98 and 0.99 (Fig. 3). In addition, TaqMan qPCR assays were performed independently for each target to determine if the primer and probe sets were as robust and accurate as in multiplex settings. For *tom*A*, rhu*M, and tomato *tubulin alpha-3*, an efficiency of 90.9%, 95.3% and 95.2 %, respectively was reported in the single qPCR assay (Fig. 3). Overall, the TaqMan qPCR multiplex is as sensitive and robust as the single TaqMan qPCR assay.

### Validation of Multiplex TaqMan qPCR with plants and seeds samples

The multiplex TaqMan qPCR was validated by analyzing DNA from tomato plants artificially inoculated with *Cm* and healthy tomato plants. *Cm* was detected in the genomic DNA from all stems, leaves, and root tissue samples of inoculated plants, whereas it was not detected in healthy plants (Table 3). Furthermore, DNA extracted from the three different seed infestation ratios tested positive for all primers and probe sets, and the multiplex qPCR was able to detect an infection level of 0.0625% (Table 4). The internal control tomato *tubulin alpha-3* amplified in all the samples except NTC (Table 3 and Table 4). No amplification was also detected when genomic DNA from healthy tomato plants and non-infected seeds was used except for the internal control tomato *tubulin alpha 3* (Tables 3 and 4).

**Table 3:** Quantification of *Clavibacter michiganensis* in inoculated M82 tomato plants using the TaqMan multiplex qPCR designed in this study in different plant tissues.

| Plant | Plant tissues | Mean C<sub>T</sub> value $\pm \sigma$ | | | Multiplex<br>gold standard |
| --- | --- | --- | --- | --- | --- |
|  |  | *tomA* (FAM) | *rhuM* (HEX) | *α-tubulin* (Cy5) |  |
| Infected 1 | Roots | 22.26 ± 1.56 | 25.11 ± 1.59 | 34.48 ± 1.53 | + |
| | Inoculation site | 17.71 ± 0.10 | 20.75 ± 0.02 | 31.53 ± 0.03 | + |
| | Stem – 3 cm | 17.12 ± 0.06 | 20.90 ± 0.09 | 32.10 ± 0.09 | + |
| | Stem – 6 cm | 18.20 ± 0.06 | 21.66 ± 0.07 | 31.38 ± 0.04 | + |
| | Leaf - Rachis | 17.56 ± 0.03 | 20.83 ± 0.05 | 33.99 ± 0.11 | + |
| Infected 2 | Roots | 21.57 ± 0.36 | 24.47 ± 0.40 | 32.60 ± 0.27 | + |
| | Inoculation site | 17.44 ± 0.03 | 20.45 ± 0.05 | 31.46 ± 0.07 | + |
| | Stem – 3 cm | 17.45 ± 0.05 | 21.23 ± 0.06 | 31.21 ± 0.02 | + |
| | Stem – 6 cm | 18.32 ± 0.09 | 22.13 ± 0.07 | 31.13 ± 0.09 | + |
| | Leaf - Rachis | 19.84 ± 0.16 | 23.24 ± 0.16 | 33.69 ± 0.11 | + |
| Infected 3 | Roots | 22.55 ± 0.16 | 25.75 ± 0.13 | 31.59 ± 0.19 | + |
| | Inoculation site | 17.56 ± 0.07 | 20.70 ± 0.12 | 31.25 ± 0.12 | + |
| | Stem – 3 cm | 19.86 ± 0.36 | 23.32 ± 0.34 | 32.16 ± 0.32 | + |
| | Stem – 6 cm | 17.57 ± 0.16 | 21.26 ± 0.18 | 30.76 ± 0.26 | + |
| | Leaf - Rachis | 19.42 ± 0.08 | 22.81 ± 0.13 | 33.95 ± 0.13 | + |
| Infected 4 | Roots | 24.35 ± 0.06 | 27.64 ± 0.17 | 31.67 ± 0.09 | + |
| | Inoculation site | 19.10 ± 0.12 | 22.38 ± 0.16 | 31.01 ± 0.07 | + |
| | Stem – 3 cm | 28.95 ± 0.10 | 32.25 ± 0.43 | 32.10 ± 0.11 | - |
| | Stem – 6 cm | 31.87 ± 0.31 | 34.99 ± 0.13 | 31.06 ± 0.17 | - |
| | Leaf - Rachis | 35.34 ± 0.11 | 36.84 ± 0.59 | 33.41 ± 0.11 | - |
| Infected 5 | Roots | 22.63 ± 0.03 | 25.90 ± 0.02 | 32.66 ± 0.07 | + |
| | Inoculation site | 19.01 ± 0.07 | 21.87 ± 0.05 | 31.09 ± 0.08 | + |
| | Stem – 3 cm | 17.23 ± 0.08 | 21.28 ± 0.09 | 30.74 ± 0.14 | + |
| | Stem – 6 cm | 18.59 ± 0.03 | 22.46 ± 0.03 | 31.09 ± 0.09 | + |
| | Leaf - Rachis | 18.57 ± 0.09 | 22.26 ± 0.07 | 32.93 ± 0.17 | + |
| Infected 6 | Roots | 20.49 ± 0.01 | 23.30 ± 0.05 | 32.23 ± 0.06 | + |
| | Inoculation site | 18.00 ± 0.05 | 21.47 ± 0.07 | 30.20 ± 0.01 | + |
| | Stem – 3 cm | 17.78 ± 0.18 | 21.30 ± 0.16 | 31.20 ± 0.20 | + |
|  | Stem – 6 cm | 18.09 ± 0.09 | 22.23 ± 0.09 | 30.60 ± 0.08 | + |
|  | Leaf - Rachis | 20.90 ± 0.13 | 24.23 ± 0.08 | 33.47 ± 0.11 | + |
| Infected 7 | Roots | 21.77 ± 0.09 | 24.91 ± 0.13 | 32.50 ± 0.10 | + |
|  | Inoculation site | 17.04 ± 0.07 | 20.63 ± 0.07 | 30.15 ± 0.09 | + |
|  | Stem – 3 cm | 19.41 ± 0.11 | 23.02 ± 0.22 | 32.08 ± 0.03 | + |
|  | Stem – 6 cm | 19.65 ± 0.07 | 23.30 ± 0.07 | 31.34 ± 0.11 | + |
|  | Leaf - Rachis | 21.99 ± 0.15 | 25.54 ± 0.21 | 33.27 ± 0.18 | + |
| Infected 8 | Roots | 21.33 ± 0.04 | 24.44 ± 0.11 | 31.79 ± 0.13 | + |
|  | Inoculation site | 18.66 ± 0.04 | 21.80 ± 0.11 | 31.10 ± 0.02 | + |
|  | Stem – 3 cm | 18.14 ± 0.04 | 21.68 ± 0.07 | 31.34 ± 0.02 | + |
|  | Stem – 6 cm | 18.03 ± 0.33 | 21.84 ± 0.24 | 30.22 ± 0.39 | + |
|  | Leaf - Rachis | 19.03 ± 0.03 | 22.26 ± 0.08 | 31.98 ± 0.07 | + |
| Infected 9 | Roots | 21.89 ± 0.06 | 24.90 ± 0.04 | 31.86 ± 0.12 | + |
|  | Inoculation site | 18.58 ± 0.14 | 21.58 ± 0.17 | 30.93 ± 0.20 | + |
|  | Stem – 3 cm | 18.36 ± 0.44 | 21.52 ± 0.36 | 30.98 ± 0.51 | + |
|  | Stem – 6 cm | 21.10 ± 0.04 | 24.45 ± 0.06 | 30.94 ± 0.03 | + |
|  | Leaf - Rachis | 23.28 ± 0.03 | 25.79 ± 0.06 | 33.18 ± 0.12 | + |
| Infected 10 | Roots | 21.13 ± 0.13 | 24.21 ± 0.12 | 31.67 ± 0.13 | + |
|  | Inoculation site | 18.15 ± 0.09 | 21.18 ± 0.06 | 31.15 ± 0.07 | + |
|  | Stem – 3 cm | 17.42 ± 0.18 | 21.04 ± 0.09 | 30.94 ± 0.20 | + |
|  | Stem – 6 cm | 18.16 ± 0.08 | 21.65 ± 0.11 | 31.02 ± 0.16 | + |
|  | Leaf - Rachis | 20.21 ± 0.09 | 23.49 ± 0.05 | 33.47 ± 0.39 | + |
| Infected 11 | Roots | 21.89 ± 0.07 | 25.03 ± 0.09 | 32.08 ± 0.13 | + |
|  | Inoculation site | 17.49 ± 0.12 | 20.80 ± 0.19 | 31.12 ± 0.09 | + |
|  | Stem – 3 cm | 18.55 ± 0.10 | 22.23 ± 0.03 | 31.96 ± 0.05 | + |
|  | Stem – 6 cm | 18.54 ± 0.12 | 22.19 ± 0.14 | 30.97 ± 0.13 | + |
|  | Leaf - Rachis | 18.61 ± 0.07 | 22.21 ± 0.08 | 33.13 ± 0.04 | + |
| Infected 12 | Roots | 21.06 ± 0.07 | 24.09 ± 0.16 | 31.72 ± 0.10 | + |
|  | Inoculation site | 18.07 ± 0.03 | 21.08 ± 0.05 | 31.46 ± 0.05 | + |
|  | Stem – 3 cm | 17.97 ± 0.41 | 21.67 ± 0.16 | 31.45 ± 0.35 | + |
|  | Stem – 6 cm | 19.57 ± 0.11 | 23.06 ± 0.11 | 31.21 ± 0.13 | + |
|  | Leaf - Rachis | 22.97 ± 0.12 | 26.25 ± 0.13 | 33.92 ± 0.03 | + |
| Healthy 1 | Roots | - | - | 31.66 ± 0.16 | - |
| | Inoculation site | - | - | $31.14 \pm 0.04$ | - |
| | Stem – 3 cm | - | - | $30.75 \pm 0.20$ | - |
| | Stem – 6 cm | - | - | $31.02 \pm 0.13$ | - |
| | Leaf - Rachis | - | - | $33.47 \pm 0.30$ | - |
| Healthy 2 | Roots | - | - | $31.70 \pm 0.08$ | - |
| | Inoculation site | - | - | $31.03 \pm 0.11$ | - |
| | Stem – 3 cm | - | - | $32.10 \pm 0.10$ | - |
| | Stem – 6 cm | - | - | $31.10 \pm 0.18$ | - |
| | Leaf - Rachis | - | - | $33.40 \pm 0.15$ | - |
| Healthy 3 | Roots | - | - | $33.28 \pm 0.089$ | - |
| | Inoculation site | - | - | $31.44 \pm 0.062$ | - |
| | Stem – 3 cm | - | - | $31.90 \pm 0.043$ | - |
| | Stem – 6 cm | - | - | $29.80 \pm 0.151$ | - |
| | Leaf - Rachis | - | - | $34.46 \pm 0.074$ | - |

$\sigma$ , standard deviation of cycle threshold (Ct) derived from the three replicates of each sample. Dash, no target amplification, negative result.

**Table 4:** Quantification of *Clavibacter michiganensis* in inoculated M82 tomato seeds by qPCR with distinct infestation levels.

| Infestation level | Mean C<sub>T</sub> value ± $\sigma$ | | |
| --- | --- | --- | --- |
|  | *tomA* (FAM) | *rhuM* (HEX) | *α-tubulin* (Cy5) |
| 100% | 16.00 ± 0.092 | 16.32 ± 0.089 | 32.12 ± 0.023 |
| 0.25% | 21.94 ± 0.032 | 22.27 ± 0.030 | 29.98 ± 0.037 |
| 0.125% | 22.29 ± 0.066 | 22.83 ± 0.096 | 29.21 ± 0.099 |
| 0.0625% | 22.42 ± 0.063 | 24.71 ± 0.183 | 28.55 ± 0.045 |
| 0% | - | - | 30.95 ± 0.076 |

- 2 $\sigma$ , standard deviation of cycle threshold (Ct) derived from the three replicates. Dash, no target amplification, negative result.

These results confirm the high specificity and accuracy of the TaqMan multiplex qPCR for the detection of *Cm* in infected plant samples as well as in infected seed samples. All the infected samples analyzed were also positive with the multiplex developed by Thapa et al. (2020) (Table 3, Fig. 4A, B). This observation was confirmed through isolation of *C. michiganensis* and the confirmation of the positive colonies through mPCR (Fig. 4B). However, while we detected *Cm* in all three different seed infestation ratios using the TaqMan multiplex qPCR, no amplification was observed for the infestation level 0.125% and 0.0625% using Thapa et al. (2020) multiplex (Fig. S1).

**Figure 4.**
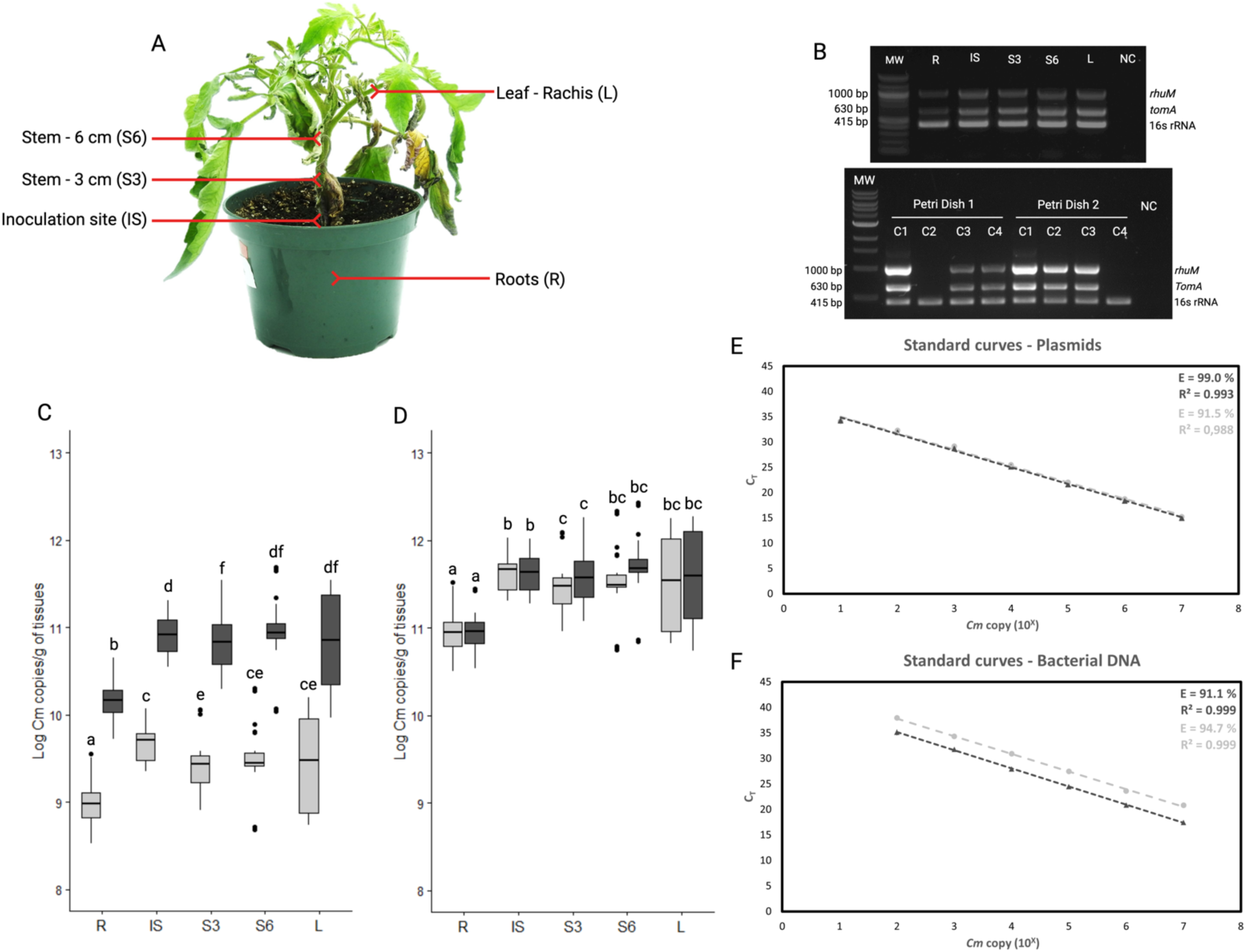
Detection of *Clavibacter michiganensis* by multiplex TaqMan qPCR in Cmm-1375 inoculated tomato plants at 21 dpi. A, parts of the plant collected for the analyses. B, Detection of *Cm* by multiplex PCR developed by Thapa et al. (2020) in the different parts of the plant. Confirmation of the isolation of *Cm* from the infected roots through colony PCR (C1-4 from two petri dishes from two infected tomato plants roots). Agarose gel electrophoresis, MW, molecular weight marker 1 kb DNA ladder (ThermoFisher, CAD). C, Number of copies/g of plant tissue obtained with the multiplex TaqMan qPCR when quantified with *tom*A (dark gray) and *rhu*M gene (light gray) plasmid-based standard curves. Different letters represent the statistically significant differences (p < 0.05) in *Cm* copies per g of tissue, according to Tukey’s HSD. D, Number of copies per g of tissue obtained with the multiplex TaqMan qPCR when quantified with *tom*A (dark gray) and *rhu*M gene (light gray) bacterial DNA-based standard curves. Statistic differences are presented as mentioned above. E, standard curves for multiplex TaqMan qPCR generated using seven-fold serial dilutions of plasmid DNA. F, standard curves for multiplex TaqMan qPCR generated using seven-fold serial dilutions of bacterial DNA. For E and F, the number of copies ranged from 10^7^ to 10^1^ copies. Dot/light gray represents *rhu*M gene, triangle/dark gray represents *tom*A gene. The values represent the mean Ct values of three technical replicates.

To quantify *Cm* in the different parts of the plants 21 dpi (Fig. 4A), *tom*A and *rhu*M copies detected in the tissues via TaqMan multiplex qPCR were transformed in absolute gene copy numbers to represent the number of bacteria per gram of tissue. For that, we used plasmid-based (Fig 4C, F) and bacterial-based standard curves (Fig. 4D, F). Using both standard curves we found that the number of copies of *Cm* in the roots was significantly lower (p < 0.05) than in other tissues (Fig. 4C-D). Interestingly, the number of copies of *Cm* is significatively lower (p < 0.05) when quantifying with *rhu*M and using the plasmid-based standard curve (Fig. 4E).

## DISCUSSION

In this study, we developed and validated a sensitive and specific TaqMan multiplex qPCR assay for the detection and quantification of *Clavibacter michiganensis*, the causal agent of bacterial canker of tomato, in plant tissue and seeds using the pathogen genes *tom*A and *rhu*M, and tomato *tubulin alpha-3*. These genes *tom*A and *rhu*M have been identified in regions conserved within *Cm* genomes and absent in other *Clavibacter* ssp. and tomato-associated bacteria species through comparative genomics (Thapa et al. 2020), pointing that they might be related to pathogenicity and/ or host specificity. These genes were thus the perfect target to develop quantitative-specific diagnostic assays aimed at the fast and accurate detection of *Cm* as previously proved through multiplex PCR (Thapa et al. 2020).

The Gram-positive genus *Clavibacter* is currently classified into nine species: *Clavibacter michiganensis*, *C. nebraskensis*, *C. capsici*, *C. sepedonicus*, *C. tessellarius*, *C. insidiosus*, *C. zhangzhiyongii*, *C. californiensis*, and *C. phaseoli* (Arizala et al. 2022). In our study, after evaluating six of these species, we have confirmed that our assay is specific to *C. michiganensis*. Furthermore, for the first time in contrast to earlier studies that developed quantitative assays for any *Clavibacter* species, we have enhanced the accuracy of our test by evaluating the specificity of the developed primers and probes against endophytic *Clavibacter* strains, thereby reducing the likelihood of false-positive results (Ouyang et al. 2013; Han et al. 2018; Ciampi-Guillardi et al. 2020; Ramachandran et al. 2021; Duong et al. 2022).

Multiplexing a real-time PCR procedure confers the advantage of detection and quantification of multiple targets in a single reaction. Thus, by adding an internal control and using two *Cm* target genes we improved the assay’s reliability and reduced the possibility of detecting false negatives. In addition, multiplexing can help to reduce the cost of the assay and reduce time because it increases confidence in the results (Arif et al. 2015; Motyka et al. 2017).

In Canada, most *Cm* outbreaks experienced by the greenhouse industry are related to contaminated seed lots that have escaped routine tests or that have not been tested at all for the presence of the pathogen (Sabaratnam 2021). Today we know that healthy seeds effectively reduce the spread of pathogens (Choudhury et al. 2017; Guimarães et al. 2017). Our multiplex TaqMan qPCR detected an infection level of 0.0625%, representing one infected seed in 1,600 seeds, a significantly higher limit of detection than the multiplex PCR used as a reference to develop our quantitative assay (Thapa et al. 2020) and higher than other qPCR assays previously developed (Ouyang et al. 2013; Han et al. 2018; Ciampi-Guillardi et al. 2020; Ramachandran et al. 2021; Duong et al. 2022). In addition, the Taqman multiplex qPCR allowed the detection of up to five copies of *Cm* genome, which is comparable or better than other qPCR developed for plant pathogenic bacteria like *Xyllela fastidiosa* (Harper et al. 2010), ‘*Candidatus* Phytoplasma asteris’ (Pérez-López et al. 2017; Hammond et al. 2021), and ‘*Candidatus* Phytoplasma rubi’ (Bennypaul et al. 2023), among others. The assay also allowed the detection of *Cm* in both plant tissue and seeds, increasing its versatility and applicability in comparison with other TaqMan multiplex qPCRs recently developed to detect and quantify *Cm* (Peňázová et al. 2020; Ramachandran et al. 2021).

The presence of *Cm* in the roots has been a subject of debate (Xu et al. 2012; Lelis et al. 2014). Here we detected, through multiplex PCR and TaqMan qPCR the presence of the bacteria in all infected plants and plant tissues analyzed, results also confirmed by isolation of the bacteria from this tissue (Table 4, Fig. 4). In plants, the amount of *Cm* was lower in the roots than in the aerial parts (Fig. 4). Naturally, *Cm* can colonize tomato plants xylem in both acropetal and basipetal directions (Xu et al. 2010, Xu et al. 2012, Chalupowicz et al. 2012, Lelis et al., 2014). Xylem sap circulates acropetally using the negative tension generated by transpiration water loss and positive pressure generated by water uptake in the roots (Venturas et al. 2017). Xylem circulation could facilitate the migration of bacteria from the roots to the shoots and increase the presence of bacteria in the apex. However, Xu et al. (2012) suggested that *Cm* moved faster to the roots than to the plant apex, finding at 20 dpi through bioluminescence, a high concentration of bacteria in the roots, stem, and leaf rachis (Xu et al. 2012), while another study found similar amounts of bacteria in all tissues (Lelis et al. 2014). It is difficult to compare the results obtained in other studies because the virulence of the isolate and the cultivar used may influence the colonization of *Cm* in the plant (Peritore-Galve et al. 2020).

One interesting finding was that when using the *tom*A and *rhu*M with the plasmid-based standard curve, there was a significant difference between the amount of *Cm* quantified with the two genes (Fig. 4C), but this difference disappeared when we quantified using a bacterial DNA-based standard curve (Fig. 4D). When using the bacterial DNA-based standard curve, there was a difference of about three cycles of amplification between the standard curve of the *tom*A and *rhu*M gene (Fig. 4F). This significant difference can be explained by the presence of multiple copies of the *tom*A gene in the strain Cmm-1375 used as positive control and to generate the standard curve. No study has shown the presence of two copies of this gene in *Cm*, and no evidence of this was found in any genome available in the NCBI. Another explanation could be that we overestimated the number of bacteria with incomplete plasmid linearization of the *rhu*M plasmid standard curve. There is evidence that supercoiled plasmid standard as a template in qPCR assays can lead to an overestimation of around eight-fold during amplification (Hou et al. 2010), the difference observed between DNA-based standard curves and the plasmid-based standard curve (Fig. 4F).

Over five million hectares of land around the world produce 189 million tons of tomato fruit per year (FAO 2021). The yield of the tomato industry can be greatly affected by bacterial canker jeopardizing food security. The TaqMan multiplex qPCR diagnostic developed here and its fast implementation in diagnostic laboratories can contribute to curbing *Cm* future outbreaks and the economic losses related to the disease. In addition, this diagnostic tool can be adopted by seed certification programs in North America and worldwide and by inspection agencies trying to stop the spread of *Cm*.

## ACKNOWLEDGEMENTS

The authors would like to thank Prof. Gitta Coaker from UC Davis, Dr. Mohammad Arif from University of Hawaii, Dr. Alejandra Huerta from North Carolina State University, and Dr. James Tambong from AAFC for providing genomic DNA of *Clavibacter spp.* strains used in this study to validate the assay. The authors would like to thank MAPAQ for supporting this work through the *Chaire de recherche en phytoprotection serricole* MAPAQ-Premier Tech, project PPIA20, and NSERC for supporting ASB through the USRA and Master’s scholarships and EPL through the Discovery program.

## SUPPLEMENTARY

**Table S1.**
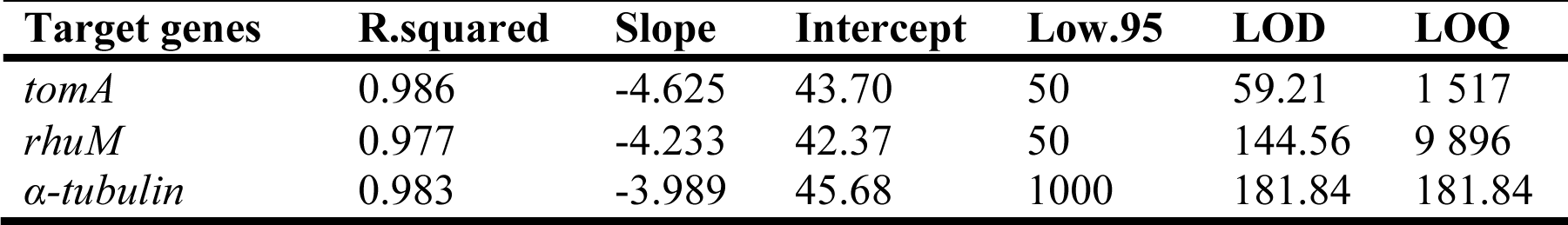
Limit of detection (LOD) and limit of quantification (LOQ) for the multiplex TaqMan qPCR assay, as determined using an optimized curve-fitting model.

**Figure S1.**
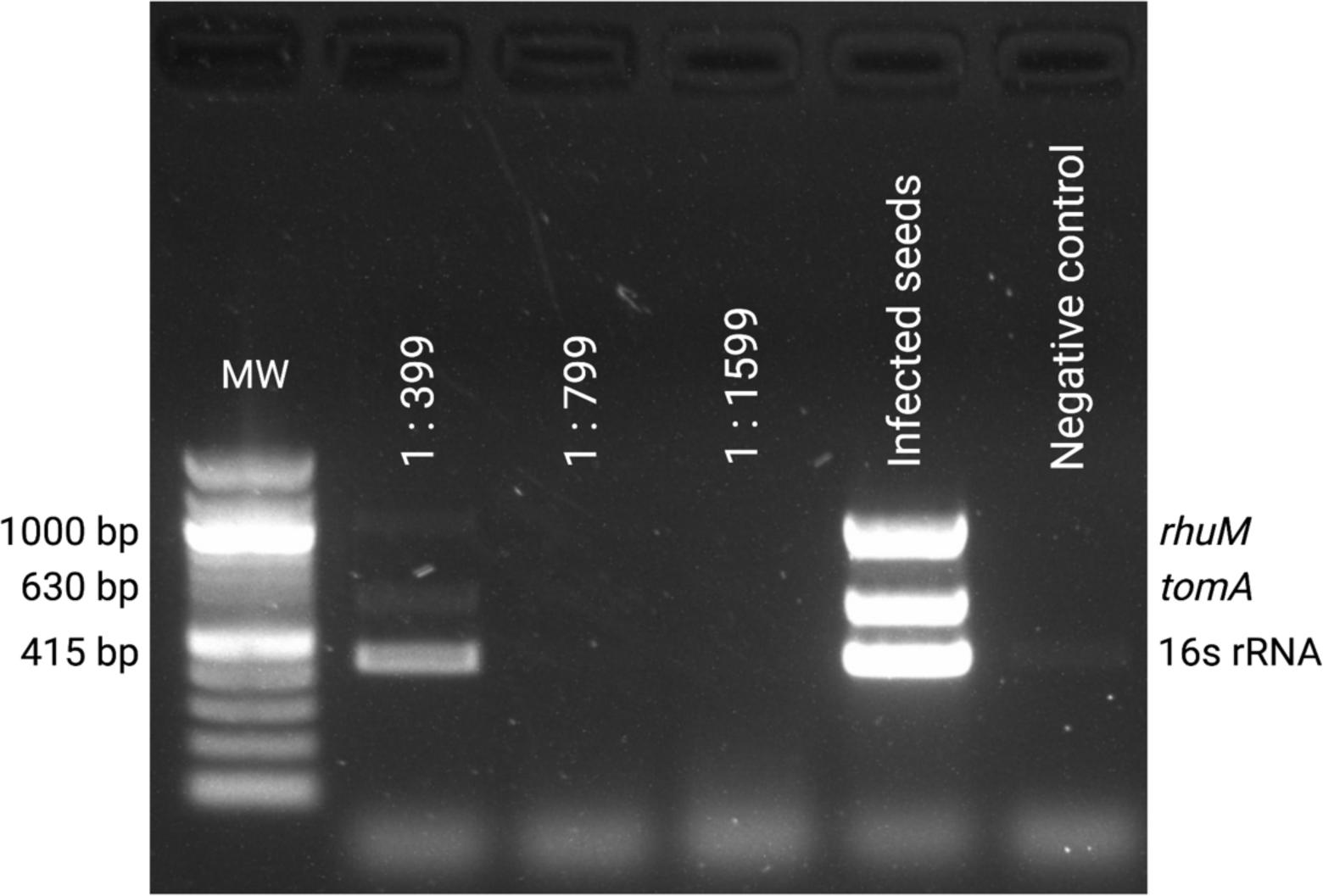
Detection from infected seeds with the multiplex developed by Thapa et al (2020). The following ratios of infested/healthy seeds were tested 1:399, 1:799, and 1:1,599, which corresponds to infestation levels of 0.25, 0.125, and 0.0625% respectively. The DNA tested in ‘infected seeds’ lane was undiluted. Agarose gel electrophoresis, MW, molecular weight marker 100 bp DNA.

### Detailed seed infection protocol and DNA extraction for *C. michiganensis* detection

*Adapted from International Seed Federation, Detection of Clavibacter michiganensis subsp. michiganensis in Tomato seed*.

### Materials

-Seed extraction buffer (Table 1.1)

-Refrigerator at 4-7 °C

-15/50 mL conical tubes

-1.5/2ml microtubes

-Centrifuge capable of spinning 15/50 mL conical tubes and 1.5/2ml microtubes

-High Speed homogenizer

-Ceramic beads (3 mm)

-Heating bath

-Vortex

-CTAB buffer (Table 1.2)

*Should be prepared on the same day.

-Ethanol 70% (high strength) ice

*Should be prepared on the same day.

-Isopropanol (2-propanol) pure ice

-Chloroform: Isoamyl alcohol (24:1)

*Can be made in advance and stored in the fridge with parafilm.

### Steps

1. Put every subsample individually into a conical tube and add sterile seed extraction buffer to each tube at a ratio of 4 mL of seed extraction buffer to 1 g of seed (v/w).
2. Incubate overnight (minimum 14 hours) at 4-7 °C.
3. Macerate for 30 min in a heat bath at 65 °C *Mix the tubes every 5 min with vortex.
4. Transfer all the fluid of extract to a suitable centrifuge tube (Tube for bead mill).
5. Centrifuge @ 14,000 g for 3 minutes.
6. Remove the supernatant carefully.
7. Add 2 ceramic beads dia. 3 mm and 400 µl of CTAB buffer heated to 60-65 °C.
8. Grind with the Omni grinder under the following conditions. S = 5 m/s T = 2 x 30 s Pause = 30 seconds
9. Add 300 µl of CTAB buffer and vortex well + mini spin down.
10. Place the tubes in 65 °C water bath for 30 minutes.
11. Centrifuge @ 14,000 g for 10 minutes.
12. Take approximately 700 µl o the supernatant and transfer to clean tubes.
13. Add an equal volume (700 µl) of Chloroform: Isoamyl alcohol (24:1) mix well (vortex).
14. Centrifuge @ 14,000 g for 10 minutes.
15. Transfer the upper fraction (water-soluble) to a new tube (approximately 550 µl).
16. Add 0.7 volume of cold isopropanol (385 µl of isopropanol for 550 µl of buffer). Mix well by inverting the tube 10 times.
17. Incubate in cold at -20 °C overnight.
18. Centrifuge the samples at 14,000 g for 15 minutes at 4 °C. Remove the supernatant (isopropanol) by decanting slowly: being careful not to break up the DNA pellet.
19. Slowly add 500 µl of ice-cold 70% EtOH to the side of the tube which is opposite to the base (in fact, it is essential at this stage not to detach the pellet from the bottom of the tube as it may get lost in the next decantation).
20. Centrifuge the samples at 14,000 g for 15 mins at 4 °C.
21. Discard the Ethanol by decantation.
22. Repeat step 19 to 21.
23. Place the tubes upside down on brown paper so that the residual ethanol is dried by capillary action for no more than 10 minutes but dried well.
24. Add 20 µl of ddH_2_O to the tubes and mix well to dissolve the DNA.
25. Quantify and determine the purity using Nanodrop or Qubit.

**Table 1.1:** Seed extraction buffer (pH 7.4)^a^.

| Compound | Amount/L |
| --- | --- |
| Na<sub>2</sub>HPO<sub>4</sub> | 7.75 g |
| KH<sub>2</sub>PO<sub>4</sub> | 1.65 g |
| Tween 20 | 0.50 mL |

<sup>a</sup>Use de-ionized water, and autoclave at 121 °C, 15 psi for 15 min.

**Table 1.2:** CETAB Buffer (CTAB 2%, Tris-HCl 100 mM, EDTA 20 mM, NaCl 1.4M)

| Compound | Amount/50 mL | Amount/25 mL |
| --- | --- | --- |
| CTAB 10% @ 37 °C | 10 mL | 5 mL |
| Tris-HCl 1 M | 5 mL | 2.5 mL |
| EDTA 0.5 M | 2 mL | 1 mL |
| NaCl 5M | 14 mL | 7 mL |
| PVP 40 | 0.5 g | 0.25 g |
| ddH2O | Volume up to 50 mL | Volume up to 25 mL |

## REFERENCES

Alvarez, A. M., Kaneshiro, W. S., and Vine, B. G. 2005. Diversity of Clavibacter michiganensis subsp. michiganensis populations in tomato seed: What is the significance? Acta Hortic. 695:205–214.

Arif, M., Opit, G., Mendoza-Yerbafría, A., Dobhal, S., Li, Z., Kučerová, Z., and Ochoa-Corona, F. M. 2015. Array of Synthetic Oligonucleotides to Generate Unique Multi-Target Artificial Positive Controls and Molecular Probe-Based Discrimination of Liposcelis Species. PLOS ONE, 10(6), e0129810.

Bennypaul, H. S., Sanderson, D., Donaghy, P., and Abdullahi, I. 2023. Development of a real-time PCR assay for the detection and identification of Rubus Stunt Phytoplasma in Rubus spp. Plant Disease, PDIS-09–22-2193-RE.

Bertelloni, F., Tosi, G., Massi, P., Fiorentini, L., Parigi, M., Cerri, D., and Ebani, V. V. 2017. Some pathogenic characters of paratyphoid Salmonella enterica strains isolated from poultry. Asian Pacific Journal of Tropical Medicine, 10(12).

Blanc-Potard, A.-B., Solomon, F., Kayser, J., and Groisman, E. A. 1999. The SPI-3 Pathogenicity Island of Salmonella enterica. Journal of Bacteriology, 181(3), 998–1004.

Chalupowicz, L., Barash, I., Reuven, M., Dror, O., Sharabani, G., Gartemann, K.-H., Eichenlaub, R., Sessa, G., and Manulis-Sasson, S. 2017. Differential contribution of Clavibacter michiganensis ssp. michiganensis virulence factors to systemic and local infection in tomato: Virulence of Cmm in systemic and local infections. Molecular Plant Pathology, 18(3), 336–346.

Chalupowicz, L., Zellermann, E.-M., Fluegel, M., Dror, O., Eichenlaub, R., Gartemann, K.-H., Savidor, A., Sessa, G., Iraki, N., Barash, I., and Manulis-Sasson, S. 2012. Colonization and Movement of GFP-Labeled Clavibacter michiganensis subsp. Michiganensis During Tomato Infection. Phytopathology, 102(1), 23–31.

Chang, R. J., Ries, S. M., and Pataky, J. K. 1991. Dissemination of Clavibacter michiganensis subsp. michiganensis by practices used to produce tomato transplants. Phytopathology. 81:1276–1281

Choudhury, R. A., Garrett, K. A., Klosterman, S. J., Subbarao, K. V., and McRoberts, N. 2017. A Framework for Optimizing Phytosanitary Thresholds in Seed Systems. Phytopathology, 107(10), 1219–1228

Ciampi-Guillardi, M., Ramiro, J., Moraes, M. H. D. de, Barbieri, M. C. G., and Massola, N. S. 2020. Multiplex qPCR Assay for Direct Detection and Quantification of Colletotrichum truncatum, Corynespora cassiicola, and Sclerotinia sclerotiorum in Soybean Seeds. Plant Disease, 104(11), 3002–3009.

Coker, J. S., and Davies, E. 2003. Selection of candidate housekeeping controls in tomato plants using EST data. BioTechniques, 35(4), 740–748.

Croce, V., Pianzzola, M. J., Durand, K., González-Arcos, M., Jacques, M.-A., and Siri, M. I. 2016. Multilocus Sequence Typing reveals high variability among *Clavibacter michiganensis* subsp. *Michiganensis* strains affecting tomato crops in Uruguay. European Journal of Plant Pathology, 144(1), 1–13.

Davis, M. J., Gillaspie, A. G., Vidaver, A. K., and Harris, R. W. 1984. Clavibacter: A New Genus Containing Some Phytopathogenic Coryneform Bacteria, Including Clavibacter xyli subsp. xyli sp. nov., subsp. nov. and Clavibacter xyli subsp. cynodontis subsp. nov., Pathogens That Cause Ratoon Stunting Disease of Sugarcane and Bermudagrass Stunting Disease. International Journal of Systematic Bacteriology, 34(2), 107–117.

de León, L., Siverio, F., and Rodríguez, A. 2006. Detection of Clavibacter michiganensis subsp. Michiganensis in tomato seeds using immunomagnetic separation. Journal of Microbiological Methods, 67(1), 141–149.

Deer, D. M., Lampel, K. A., and González-Escalona, N. 2010. A versatile internal control for use as DNA in real-time PCR and as RNA in real-time reverse transcription PCR assays. Letters in Applied Microbiology, 50(4), 366–372.

Dreier, J., Meletzus, D., and Eichenlaub, R. 1997. Characterization of the Plasmid Encoded Virulence Region pat -1 of Phytopathogenic Clavibacter michiganensis subsp. Michiganensis. Molecular Plant-Microbe Interactions, 10(2), 195–206.

Duong, H. T., Williams, B., White, D., Burgess, T. I., and Hardy, G. E. St. J. 2022. QPCR Assays for Sensitive and Rapid Detection of Quambalaria Species from Plant Tissues. Plant Disease, 106(1), 107–113.

Ebner, P., and Götz, F. 2019. Bacterial Excretion of Cytoplasmic Proteins (ECP): Occurrence, Mechanism, and Function. Trends in Microbiology, 27(2), 176–187

EPPO. 2016. PM 7/42 (3) *Clavibacter michiganensis* subsp. *michiganensis*. EPPO Bulletin, 46(2), 202–225.

FAO. 2019. FAOSTAT. Food and Agriculture Data. www.FAO.org/faostat/

Fatmi, M. B., Bolkan, H., Schaad, N. W., Walcott, R. R., and Schaad, N. W. 2017. Detection of *Clavibacter michiganensis* subsp. *michiganensis* in tomato seeds. Detection of Plant Pathogenic Bacteria in Seed and Other Planting Material, 111-7.

Ftayeh, R. M., von Tiedemann, A., and Rudolph, K. W. E. 2011. A New Selective Medium for Isolation of *Clavibacter michiganensis* subsp. *Michiganensis* from Tomato Plants and Seed. Phytopathology, 101: 1355–1364.

Franken, A. A. J. M., Kamminga, G. C., Snijders, W., Zouwen, P. S., and Birnbaum, Y. E. 1993. Detection of *Clavibacter michiganensis* ssp. *Michiganensis* in tomato seeds by immunofluorescence microscopy and dilution plating. Netherlands Journal of Plant Pathology, 99(3), 125–137.

Friedman, M. (2002). Tomato Glycoalkaloids: Role in the Plant and in the Diet. Journal of Agricultural and Food Chemistry, 50(21), 5751–5780.

Gartemann, K.-H., Abt, B., Bekel, T., Burger, A., Engemann, J., Flügel, M., Gaigalat, L., Goesmann, A., Gräfen, I., Kalinowski, J., Kaup, O., Kirchner, O., Krause, L., Linke, B., McHardy, A., Meyer, F., Pohle, S., Rückert, C., Schneiker, S., … Bartels, D. 2008. The Genome Sequence of the Tomato-Pathogenic Actinomycete *Clavibacter michiganensis* subsp. Michiganensis NCPPB382 Reveals a Large Island Involved in Pathogenicity. Journal of Bacteriology, 190(6), 2138–2149.

Ghareeb, H., Bozsó, Z., Ott, P. G., and Wydra, K. 2011. Silicon and Ralstonia solanacearum modulate expression stability of housekeeping genes in tomato. Physiological and Molecular Plant Pathology, 75(4), 176–179.

Gitaiti, R. Beaver. R., Voloudakis, A. 1991. Detection of *Clavibacter michiganensis* subsp. *michiganensis* in symptomless tomato transplants. Plant Disease, 75, 834 – 8.

Guimarães, M. de R. F., Siqueira, C. da S., Machado, J. da C., França, S. K. S. de, and Guimarães, G. C. 2017. Evaluation of inoculum potential of pathogens in seeds: Relation to physiological quality and DNA quantification by *qPCR*. Journal of Seed Science, 39(3), 224–233.

Gupta, R., Lee, S. E., Agrawal, G. K., Rakwal, R., Park, S., Wang, Y., and Kim, S. T. 2015. Understanding the plant-pathogen interactions in the context of proteomics-generated apoplastic proteins inventory. Frontiers in Plant Science, 6.

Green, M. R., and Sambrook, J. 2016. Precipitation of DNA with Ethanol. Cold Spring Harbor Protocols, 2016:12.

Hadas, R., Kritzman, G., Klietman, F., Gefen, T., and Manulis, S. 2005. Comparison of extraction procedures and determination of the detection threshold for *Clavibacter michiganensis* ssp. Michiganensis in tomato seeds. Plant Pathology, 54(5), 643–649.

Hammond, C., Pérez-López, E., Town, J., Vincent, C., Moreau, D., and Dumonceaux, T. 2021. Detection of blueberry stunt phytoplasma in Eastern Canada using cpn60-based molecular diagnostic assays. Scientific Reports, 11(1), 22118.

Han, S., Jiang, N., Lv, Q., Kan, Y., Hao, J., Li, J., and Luo, L. 2018. Detection of *Clavibacter michiganensis* subsp. *michiganensis* in viable but nonculturable state from tomato seed using improved qPCR. PLoS One, 3;13(5):e0196525.

Harper, S. J., Ward, L. I., and Clover, G. R. G. 2010. Development of LAMP and Real-Time PCR Methods for the Rapid Detection of *Xylella fastidiosa* for Quarantine and Field Applications. Phytopathology, 100(12), 1282–1288.

Hou, Y., Zhang, H., Miranda, L., and Lin, S. 2010. Serious Overestimation in Quantitative PCR by Circular (Supercoiled) Plasmid Standard: Microalgal pcna as the Model Gene. PLoS ONE, 5(3)

Jacques, M.-A., Durand, K., Orgeur, G., Balidas, S., Fricot, C., Bonneau, S., Quillévéré, A., Audusseau, C., Olivier, V., Grimault, V., and Mathis, R. 2012. Phylogenetic Analysis and Polyphasic Characterization of Clavibacter michiganensis Strains Isolated from Tomato Seeds Reveal that Nonpathogenic Strains Are Distinct from *C. michiganensis* subsp. *michiganensis*. Applied and Environmental Microbiology, 78(23), 8388–8402.

Kaneshiro, W. S., Mizumoto, C. Y., and Alvarez, A. M. 2006. Differentiation of Clavibacter michiganensis subsp. Michiganensis from seed-borne saprophytes using ELISA, Biolog and 16S rDNA sequencing. European Journal of Plant Pathology, 116(1), 45–56.

Kaup, O., Gräfen, I., Zellermann, E.-M., Eichenlaub, R., and Gartemann, K.-H. 2005. Identification of a Tomatinase in the Tomato-Pathogenic Actinomycete Clavibacter michiganensis subsp. Michiganensis NCPPB382. Molecular Plant-Microbe Interactions, 18(10), 1090–1098.

Keeling, P J and Doolittle, W F. 1996. Alpha-tubulin from early-diverging eukaryotic lineages and the evolution of the tubulin family., Molecular Biology and Evolution, 13(10), 1297–1305,

Kleitman, F., Barash, I., Burger, A., Iraki, N., Falah, Y., Sessa, G., Weinthal, D., Chalupowicz, L., Gartemann, K. H., Eichenlaub, R., and Manulis-Sasson, S. 2008. Characterization of a *Clavibacter michiganensis* subsp. *michiganensis* population in Israel. Eur. J. Plant Pathol. 121:463–475.

Klymus, K. E., Merkes, C.M., Allison, M. J., Goldberg, C.S., Helbing, C.C., Hunter, M.E., Jackson, C.A., Lance, R.F., Mangan, A.M., Monroe, E.M., Piaggio, A.J., Stokdyk, J.P., Wilson, C.C., and Richter, C.A. 2019. Reporting the limits of detection and quantification for environmental DNA assays. Environ. DNA 2, 271–282.

Kokošková, B., Mráz, I., and Fousek, J. 2010. Comparison of specificity and sensitivity of immunochemical and molecular techniques for determination of *Clavibacter michiganensis* subsp. Michiganensis. Folia Microbiologica, 55(3), 239–244

Kralik, P., and Ricchi, M. 2017. A Basic Guide to Real Time PCR in Microbial Diagnostics: Definitions, Parameters, and Everything. Frontiers in Microbiology, 8.

Krämer, I., and Griesbach, E. 1995. Use of ELISA for detection of *Clavibacter michiganensis* subsp. *michiganensis* in tomato. EPPO Bulletin, 25(1–2), 185–193.

Lelis, F. M. V., Czajkowski, R., de Souza, R. M., Ribeiro, D. H., and van der Wolf, J. M. 2014. Studies on the colonization of axenically grown tomato plants by a GFP-tagged strain of *Clavibacter michiganensis* subsp. *michiganensis*. European Journal of Plant Pathology, 139(1), 53–66.

Li, X., Tambong, J., Yuan, K. (Xiaoli), Chen, W., Xu, H., Lévesque, C. A., and De Boer, S. H. 2018. Re-classification of Clavibacter michiganensis subspecies on the basis of whole-genome and multi-locus sequence analyses. International Journal of Systematic and Evolutionary Microbiology, 68(1), 234–240.

Méndez, V., Valenzuela, M., Salvà-Serra, F., Jaén-Luchoro, D., Besoain, X., Moore, E.R.B., and Seeger, M. 2020. Comparative Genomics of Pathogenic *Clavibacter michiganensis* subsp. *michiganensis* Strains from Chile Reveals Potential Virulence Features for Tomato Plants. Microorganisms, 8, 1679.

Mirmajlessi, S. M., Loit, E., Mänd, M., and Mansouripour, S. M. 2015. Real-time PCR applied to study on plant pathogens: Potential applications in diagnosis - a review. Plant Protection Science, 51(4), 177–190.

Motyka, A., Zoledowska, S., Sledz, W., and Lojkowska, E. 2017. Molecular methods as tools to control plant diseases caused by Dickeya and Pectobacterium spp: A minireview. New Biotechnology, 39, 181–189.

Murray, M. G., and Thompson, W. F. 1980. Rapid isolation of high molecular weight plant DNA. Nucleic Acids Res. 8:4321–4326.

Nandi, M., Macdonald, J., Liu, P., Weselowski, B., Yuan, ZC. 2018. *Clavibacter michiganensis* ssp. *michiganensis*: bacterial canker of tomato, molecular interactions and disease management. Mol Plant Pathol.12;19(8):2036–50.

Nakayasu, M., Ohno, K., Takamatsu, K., Aoki, Y., Yamazaki, S., Takase, H., Shoji, T., Yazaki, K., and Sugiyama, A. 2021. Tomato roots secrete tomatine to modulate the bacterial assemblage of the rhizosphere. Plant Physiology, 186(1), 270–284

Ouyang, P., Arif, M., Fletcher, J., Melcher, U., and Ochoa Corona, F. M. 2013. Enhanced Reliability and Accuracy for Field Deployable Bioforensic Detection and Discrimination of Xylella fastidiosa subsp. Pauca, Causal Agent of Citrus Variegated Chlorosis Using Razor Ex Technology and TaqMan Quantitative PCR. PLoS ONE, 8(11), e81647.

Pastrik, K. H., and Rainey, F. A. 1999. Identification and Differentiation of Clavibacter michiganensis Subspecies by Polymerase Chain Reaction-based Techniques. Journal of Phytopathology, 147(11–12), 687–693.

Peritore-Galve, F. C., Tancos, M. A., and Smart, C. D. 2021. Bacterial Canker of Tomato: Revisiting a Global and Economically Damaging Seedborne Pathogen. Plant Disease, 105(6), 1581–1595.

Peňázová E, Dvořák M, Ragasová L, Kiss T, Pečenka J, and al. 2020. Multiplex real-time PCR for the detection of Clavibacter michiganensis subsp. michiganensis, Pseudomonas syringae pv. tomato and pathogenic Xanthomonas species on tomato plants. PLOS ONE 15(1): e0227559.

Pérez-López, E., Rodríguez-Martínez, D., Olivier, C. Y., Luna-Rodríguez, M., and Dumonceaux, T. J. 2017. Molecular diagnostic assays based on cpn60 UT sequences reveal the geographic distribution of subgroup 16SrXIII-(A/I)I phytoplasma in Mexico. Scientific Reports, 7(1), 950.

Peritore-Galve, F. C., Miller, C., and Smart, C. D. 2023. Characterizing Colonization Patterns of Clavibacter michiganensis During Infection of Tolerant Wild Solanum Species. Phytopathology, 110(3), 574–581.

R Core Team. 2023. R: A language and environment for statistical computing. R Foundation for Statistical Computing, Vienna, Austria. https://www.R-project.org/

Ramachandran, S., Dobhal, S., Alvarez, A. M., and Arif, M. 2021. Improved multiplex TaqMan qPCR assay with universal internal control offers reliable and accurate detection of Clavibacter michiganensis. Journal of Applied Microbiology, 131(3)

Sabaratnam, S. 2021. Bacterial Canker of Greenhouse Tomato. Ministry of agriculture, Food and Fisheries.

Santos, M. S., Cruz, L., Norskov, P., and Rasmussen, O. F. 1997. A rapid and sensitive detection of Clavibacter michiganensis subsp. michiganensis in tomato seeds by polymerase chain reaction. Seed Sci. Technol. 25:581–584.

Sandrock, R. W., and VanEtten, H. D. 1998. Fungal Sensitivity to and Enzymatic Degradation of the Phytoanticipin α-Tomatine. Phytopathology, 88(2), 137–143.

Sen, Y., Wolf, van der, J. M., Visser, R. G. F., and Heusden, van, A. W. 2015. Bacterial canker of tomato: current knowledge of detection, management, resistance, and interactions. Plant Disease, 99(1), 4–13.

Smith, E. 1910. A new tomato disease of economic importance. Science, 31, 794–796

Tancos, M. A., Chalupowicz, L., Barash, I., Manulis-Sasson, S., and Smart, C. D. 2013. Tomato Fruit and Seed Colonization by Clavibacter michiganensis subsp. Michiganensis through External and Internal Routes. Applied and Environmental Microbiology, 79(22), 6948–6957.

Tancos, M. A., Lange, H. W., and Smart, C. D. 2015. Characterizing the Genetic Diversity of the Clavibacter michiganensis subsp. Michiganensis Population in New York. Phytopathology, 105(2), 169–179.

Tenor, J. L., McCormick, B. A., Ausubel, F. M., and Aballay, A. 2004. Caenorhabditis elegans-Based Screen Identifies Salmonella Virulence Factors Required for Conserved Host-Pathogen Interactions. Current Biology, 14(11), 1018–1024.

Thapa, S. P., O’Leary, M., Jacques, M.-A., Gilbertson, R. L., Coaker, G. 2020. Comparative Genomics to Develop a Specific Multiplex PCR Assay for Detection of *Clavibacter michiganensis*. Phytopathology, 110:556–566.

Thapa, S. P., Pattathil, S., Hahn, M. G., Jacques, M.-A., Gilbertson, R. L., and Coaker, G. 2017. Genomic Analysis of *Clavibacter michiganensis* Reveals Insight Into Virulence Strategies and Genetic Diversity of a Gram-Positive Bacterial Pathogen. Molecular Plant-Microbe Interactions, 30(10), 786–802

Venturas, M. D., Sperry, J. S., and Hacke, U. G. 2017. Plant xylem hydraulics: What we understand, current research, and future challenges. Journal of Integrative Plant Biology, 59(6), 356–389.

Wang, J., Yu, W., Yang, Y., Li, X., Chen, T., Liu, T., Ma, N., Yang, X., Liu, R., and Zhang, B. 2015. Genome-wide analysis of tomato long non-coding RNAs and identification as endogenous target mimic for microRNA in response to TYLCV infection. Scientific Reports, 5(1), 16946.

Xu, X., Miller, S. A., Baysal-Gurel, F., Gartemann, K.-H., Eichenlaub, R., and Rajashekara, G. 2010. Bioluminescence Imaging of Clavibacter michiganensis subsp. Michiganensis Infection of Tomato Seeds and Plants. Applied and Environmental Microbiology, 76(12), 3978–3988.

Xu, X., Rajashekara, G., Paul, P. A., and Miller, S. A. 2012. Colonization of Tomato Seedlings by Bioluminescent Clavibacter michiganensis subsp. Michiganensis Under Different Humidity Regimes. Phytopathology, 102(2), 177–184.

Yang, W., and Francis, D. M. 2005. Marker-assisted Selection for Combining Resistance to Bacterial Spot and Bacterial Speck in Tomato, Journal of the American Society for Horticultural Science jashs, 130(5), 716–721.

Yang, Y., Zhou, Q., Zahr, K., Harding, M. W., Feindel, D., and Feng, J. 2021. Impact of DNA extraction efficiency on the sensitivity of PCR-based plant disease diagnosis and pathogen quantification. European Journal of Plant Pathology, 159(3), 583–591.

Yasuhara-Bell, J., and Alvarez, A. M. 2015. Seed-associated subspecies of the genus Clavibacter are clearly distinguishable from Clavibacter michiganensis subsp. Michiganensis. International Journal of Systematic and Evolutionary Microbiology, 65(Pt_3), 811–826.

Zaluga, J., Van Vaerenbergh, J., Stragier, P., Maes, M., and De Vos, P. 2013. Genetic diversity of non-pathogenic Clavibacter strains isolated from tomato seeds. Systematic and Applied Microbiology, 36(6), 426–435.

Zou, W., Al-Khaldi, S. F., Branham, W. S., Han, T., Fuscoe, J. C., Han, J., Foley, S. L., Xu, J., Fang, H., Cerniglia, C. E., and Nayak, R. 2010. Microarray analysis of virulence gene profiles in Salmonella serovars from food/food animal environment. The Journal of Infection in Developing Countries, 5(02), 94–105.

